# Characterisation of the axon initial segment and intrinsic excitability in the sub-acute phase post-ischemic stroke

**DOI:** 10.1101/2025.01.09.632289

**Authors:** Emily S King, Jamie L Beros, Hakuei Fujiyama, John N.J Reynolds, Alexander D Tang

## Abstract

**Background:** Stroke is a leading cause of disability and stroke-induced changes in cortical excitability are thought to impede functional recovery. Identifying cellular targets that contribute to maladaptive excitability holds great potential for the development of therapeutic interventions to improve stroke outcomes. One potential target is the axon initial segment (AIS), the specialised cellular domain where action potentials are initiated. In the acute phase post stroke, neurons in the peri-infarct zone display abnormal AIS structural properties which is assumed to contribute to altered neuronal excitability. However, whether this continues into the sub-acute phase post stroke, a period with heightened plasticity and when physical rehabilitation typically begins is unknown.

**Methods:** We induced a photothrombotic ischemic stroke to the right motor cortex of 13-week-old mice alongside adeno-associated virus labelling of layer 2/3 and layer 5 pyramidal neurons in the peri-infarct zone and contralesional motor cortex. Immunofluorescence staining for Ankyrin-G and whole-cell patch clamp electrophysiology measures were made at 28-days post stroke to assess changes in AIS structure and function. Additionally, we investigated potential hemispheric-, cortical layer-, and sex-dependent differences in AIS and intrinsic excitability properties.

**Results:** We found that normal AIS structure and function are preserved in the sub-acute phase post ischemic stroke. However, we found evidence of reduced input resistance across both hemispheres and reduced evoked spike firing frequency in the peri-infarct zone in both sexes. In addition, we found stroke reduced the evoked spike firing frequencies in the contralesional hemisphere, but only in males.

**Conclusion:** Despite the preservation of normal AIS structure and function in the sub-acute phase post ischemic stroke, cortical pyramidal neuron excitability is reduced through other intrinsic membrane mechanisms. Additionally, we show that changes to neuronal excitability spread to the contralesional hemisphere in males. These findings provide novel insight into the maladaptive changes to neural excitability in the sub-acute phase of ischemic stroke and further highlight the need to develop sex-specific stroke treatments.

## Introduction

Stroke is a global health crisis and the third-leading cause of death and acquired disability worldwide ^1^. To date, therapies aimed at promoting recovery following stroke are limited, leaving many stroke patients with long-term disabilities and neurological deficits that impede their independence and quality of life. Post-acute restorative therapies, such as neuro-rehabilitation, remain the gold standard for reducing long-term disability in stroke patients ^2^. However, its efficacy can vary substantially due to factors such as stroke severity, prior functional status^3^, therapy consistency and intensity^2^, patient age, and sex ^4–7^. Moreover, while some stroke patients respond well to neurorehabilitation, complete restoration of function is rare and there is a critical need for more effective treatment strategies. To achieve this, a greater understanding of the pathophysiological changes that occur post stroke is needed, and how this varies with key biological factors such as sex.

Following ischemic stroke, surviving neural circuits in both cortical hemispheres show abnormal excitability (see Joy and Carmichael, 2020 for a comprehensive review) which is believed to be a barrier for functional recovery. For example, animal and clinical models of ischemic stroke report attenuated neuronal excitability in the peri-infarct zone ^9–11^, which is thought to result from an upregulation of extra-synaptic GABAergic signaling ^11^, in turn inhibiting the excitability of the pyramidal neurons in this region and the connected cortical areas in both the ipsi- and contralesional hemispheres. Additionally, acute hyperexcitability of the contralesional hemisphere has been observed in animal ^12,13^ and clinical ^9^ models of unilateral stroke, and this is thought to further inhibit excitability in the ipsilesional hemisphere (i.e. interhemispheric imbalance model of stroke; see Boddington and Reynolds, 2017 for a comprehensive review).

While altered cortical excitability following stroke is often attributed to changes at the synapse, intrinsic membrane properties outside of the synapse are also powerful regulators of neuronal excitability and have been shown to become maladaptive post ischemic stroke ^14,15^. One region where this has been shown to occur is the axon initial segment (AIS), a highly excitable axonal domain responsible for action potential (AP) generation. In the intact nervous system, the AIS undergoes dynamic periods of homeostatic plasticity in which its length, location along the axon, and/or composition and kinetics of voltage-gated ion channels are altered to fine-tune neuronal excitability and regulate AP firing properties ^16,17^. However, in the acute phase post stroke, cortical pyramidal neurons in the peri-infarct zone display altered AIS morphology, including a reduction in AIS length occurring from the distal end ^18,19^, an increase in the number of newly sprouted AISs, and a loss in the number of GABAergic axoaxonic synapses onto the AIS ^19^. These structural changes are typically associated with a reduction in neuronal excitability and provide a non-synaptic mechanism of abnormal neuronal excitability in the peri-infarct zone. As a result, the AIS has been identified as a therapeutic target for stroke treatment, but the extent of maladaptive AIS structure and function remains unclear. For example, it is unknown whether maladaptive AIS changes are restricted to the peri infract area and whether any changes last into the sub-acute phase when rehabilitation typically begins. Furthermore, sex-dependent differences in AIS plasticity and other intrinsic excitability properties post stroke, irrespective of the timepoint, remain unclear.

Here, we induced a photothrombotic ischemic stroke in the right motor cortex of 13-week-old mice and investigated changes to AIS structure and function, along with changes in intrinsic excitability 28 days later, to coincide with the sub-acute phase post stroke. Additionally, bilateral adeno-associated virus (AAV) labelling of layer 2/3 and layer 5 pyramidal neurons in the peri-infarct zone and contralesional hemisphere allowed us to investigate potential hemispheric (i.e. ipsilesional vs contralesional), cortical layer (i.e. layer 2/3 and layer 5), and sex-dependent differences in AIS and intrinsic excitability properties. Our results show that normal AIS structure and function is preserved in sub-acute phase post stroke, however, intrinsic excitability was reduced that differed between hemispheres, cortical layers, and sex. These findings highlight intrinsic excitability but not the AIS as a contributor to the dysregulation of cortical excitability observed in the sub-acute phase post stroke and supports the need to develop sex-specific treatments.

## Methods

### Animals

Animal procedures were approved by the UWA Animal Ethics Committee prior to commencement (RA/3/100/1677) and all experiments were performed in accordance with the Australian code for the care and use of animals for scientific purposes. Animals were collectively housed under an artificial 12:12h light-dark cycle, with free access to food and water. 12-week-old wildtype C57BI/6J mice of both sexes underwent one week of acclimatisation before random assignment (random animal ID number selection) to either stroke or sham/control experimental groups. Cages contained a mix of stroke and sham animals and experimenters were only blinded to the group allocation during the analysis stage. A total of 15 mice (8 stroke (3 female and 5 male) and 7 sham (4 female and 3 male)) were used to assess changes in AIS structure, while 12 mice (6 stroke and 6 sham, evenly split by sex) were used to quantify electrophysiological changes in the sub-acute phase post stroke.

### Photothrombotic stroke location and virus injection

Stereotaxic surgery was performed on all mice at 13-weeks of age to deliver injections of AAVs and the induction of a photothrombotic ischemic stroke. Buprenorphine (0.05mg/kg: intraperitoneal injection) was administered 30 minutes prior to anaesthesia with isoflurane. Once anesthetised, mice were head-fixed in a stereotaxic frame and a line block (bupivacaine, 0.2ml of 0.25mg/ml: subcutaneous) was administered to the incision site. Saline (0.01ml: intraperitoneal, every 15 minutes) was administered for hydration. A midline sagittal incision was made to expose the skull and a 0.8mm diameter craniotomy was made to the skull overlying the (left) contralesional motor cortex (1.33 mm anterior and 2 mm lateral to Bregma) and the ipsilesional (right) peri-infarct zone (0.07 mm posterior and 2 mm lateral to Bregma) for virus injections. An additional marking was made over the right forelimb region of the motor cortex (1.33 mm anterior and 2 mm lateral to Bregma) to denote the stroke site.

### Virus injections

A glass micropipette attached to a nanolitre injector (World Precision Instrument Sarasota, USA) was used to deliver bilateral virus injections to the contralesional motor cortex and the peri-infarct zone. To preferentially label excitatory neurons in the motor cortex, we used AAV5-CamKII-eGFP (Addgene, #105541-AAV5). To target the virus to layer 2/3 and layer 5 cortical motor neurons, one injection (200 nl per injection at a rate of 40nl/minute) was made at depths of 250µm and 400µm from the brain surface, respectively (Fig 1. A). Following the delivery of each injection, the micropipette was left undisturbed for one minute. Once all four injections were complete, the skin was replaced over each craniotomy in preparation for the stroke induction.

**Figure 1.**
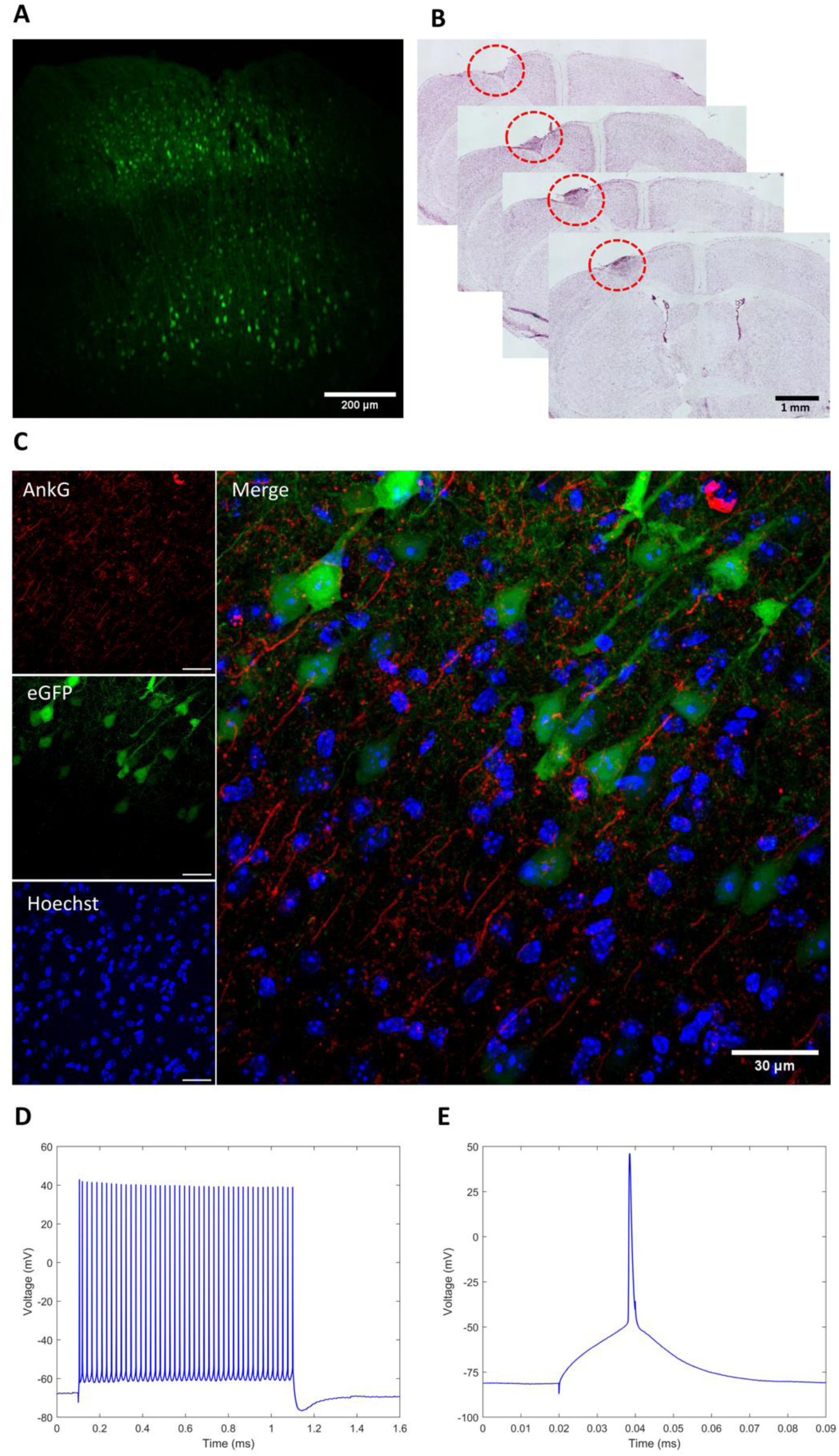
Representative images of AAV-labelling of layer 2/3 and layer 5 pyramidal neurons in the motor cortex (A), stroke lesion volume (B), maximum intensity z-projection of AAV- and immunolabelled cortical neurons and AIS used for analysis (C), and representative AP (D) and single AP (E) traces.

### Photothrombotic stroke

The stroke site was confined to the right forelimb region of the motor cortex by placing an aluminium foil sheet over the exposed skull, with a 2mm diameter cut-out made over the previously marked right motor cortex. A cold light source was then positioned directly over the 2mm cut out. The photosensitive dye Rose Bengal (Sigma Aldrich; catalogue no: 330000) was administered via intraperitoneal injection at a concentration of 15mg/Kg. 5 minutes after the Rose Bengal injection, the brain was illuminated for 15 minutes to induce an ischemic stroke (Fig 1. B), a protocol known to induce stroke lesions and functional deficits ^20^. After illumination, the incision line was sutured closed and topical anaesthetic (bupivacaine: 2.5mg/ml) was applied to the wound site. Mice were then left to recover in a dark cage for ∼ 5 hours post-surgery before being returned to their home cage. For sham control groups, mice underwent the same surgical, virus injection and light illumination procedures, but were injected with an equivalent volume of 0.9% saline instead of Rose Bengal.

### Structural Analysis

28 days following surgery, mice were deeply anesthetised with methoxyflurane (gas, 0.5ml) before lethal intraperitoneal injection of pentobarbital (intraperitoneal, 0.15ml of 160mg/kg). After confirmation of death, mice were transcardially perfused with cold 0.9% saline followed by 2% paraformaldehyde (PFA) in 0.1M PBS for 10 minutes at a flow rate of 10ml/minute. Brains were extracted and post-fixed in 2% PFA at 4°C for 45 minutes before being transferred into PBS + 0.1% sodium azide and stored at 4°C. Brains were cryoprotected in sucrose (15% in PBS for 24 hours followed by 30% in PBS for a further 24 hours), and serially sectioned in the coronal plane at a thickness of 30µm on a cryostat for either immunofluorescence or cresyl violet staining.

### Lesion volume analysis

To quantify lesion volume and position, brain sections were stained with cresyl violet using the following procedure: first, brain sections were completely immersed in distilled water for 2 minutes to wash off excess OCT, before being transferred into a solution comprising 95% ethanol and 5% acetic acid (warmed to ∼ 50°C) for 5 minutes. Sections were rinsed in distilled water for 1 minute, incubated in cresyl violet for 5 minutes, and transferred back into 95% ethanol and 5% acetic acid for 2 minutes. Sections were then dehydrated in 100% ethanol for 3 minutes, cleared in xylene for 1 minute, and covered with DPX mountant and a coverslip. Brain sections were imaged at 10x magnification on a Nikon NE-i microscope (NA = 0.3) (Fig 1. B). For volume quantification, Fiji image processing software was used manually to trace the outline of the lesion in each section. Each area was then multiplied by the thickness of the section (30um) before all area calculations were summed together and divided by 10^-9^ to obtain the lesion volume in mm^3^. To obtain a total lesion volume, the lesion volume measured in the brain sections were multiplied by 3, as every third brain section collected was dedicated to cresyl staining for lesion analysis (Supplementary Table 1).

### Immunofluorescence

For immunofluorescence staining, sections were washed with 0.1M PBS (3x 5 minutes), blocked for 3 hours at room temperature in 0.1% Triton-X (100) in 0.1M PBS + 10% normal goat serum, and then incubated for 20 hours at 4°C in 0.1M PBS containing 0.1% Triton-X, 3% normal goat serum and the primary antibody Anti-Ankyrin-G NeuroMab clone K106/36 (1:200, Developmental Studies Hydridoma Bank; catalogue no: N106/36). After primary antibody incubation, sections were washed with 0.1M PBS (3×10 minutes) and incubated for 3 hours at room temperature in the same primary blocking solution with the secondary antibody goat anti-chicken Alexa Fluor 647 (1:600. Invitrogen; catalogue no: A-21449). The nuclear label Hoechst (1:1000, Invitrogen; catalogue no: H21486) was included in the secondary antibody incubation as a non-specific cell marker. Following this, sections were washed with 0.1M PBS (3x 10 minutes), cover slipped with Prolong Diamond Antifade Mountant (Invitrogen; catalogue no: P36970) and stored at 4°C for a minimum of 48 hours prior to imaging.

### Confocal Microscopy and AIS quantification

Sections were imaged on a C2 confocal microscope (Nikon) at 60x magnification (oil immersion, NA=1.40, Nikon Plan Apo). Images were acquired in NIS Elements (Nikon) at a resolution of 512 x 512 pixels. Two fields of view were captured for each cortical layer and hemisphere for all animals. Multiple images were captured in the z-plane of each field of view, at a z-step depth of 0.25um, capturing the appearance and disappearance of all AIS fluorescence in the field of view. Images were processed in FIJI (Image J) to generate a single RGB TIFF of the maximum z projection (Fig 1 C). Prior to analysis, investigators were blinded to the condition groups (i.e. stroke vs sham control).

AIS length was quantified using a custom MATLAB code developed by Grubb and colleagues^16^. Briefly, the AIS was manually traced using the Ankyrin G fluorescence intensity, with the start and end position of the AIS defined as the first and last point along the axon where the Ankyrin G fluorescence profile diminished to 0.33 of the maximum fluorescence value. AIS distance from the soma was estimated using Fiji image processing software by measuring the distance between the end of the soma/eGFP fluorescence signal and the start of AIS/Ankyrin G fluorescence signal.

To obtain AIS structural values, AIS length and distance from the soma were averaged across cell counts to provide one measurement for each layer of each hemisphere (i.e. contralesional layer 2/3, contralesional layer 5, peri-infarct layer 2/3, peri-infarct layer 5) per animal. A total cell count across all animals is presented in Table 1.

**Table 1.** Total cell counts for layer 2/3 and layer 5 across peri-infarct and contralesional hemispheres.

|  | Stroke |  | Sham |  |
| --- | --- | --- | --- | --- |
|  | Length (μm) | Distance (μm) | Length (μm) | Distance (μm) |
| **Contralesional layer 2/3** |  |  |  |  |
| no. cells: | 590 | 82 | 497 | 56 |
| no. mice: | 8 | 7 | 6 | 5 |
| **Contralesional layer 5** |  |  |  |  |
| no. cells: | 632 | 112 | 362 | 34 |
| no. mice: | 8 | 6 | 5 | 4 |
| **Peri-infarct layer 2/3** |  |  |  |  |
| no. cells: | 324 | 26 | 222 | 24 |
| no. mice: | 5 | 3 | 4 | 3 |
| **Peri-infarct layer 5** |  |  |  |  |
| no. cells: | 501 | 83 | 191 | 35 |
| no. mice: | 7 | 7 | 3 | 2 |

### Brain slice preparation

Mice were terminally euthanised in a two-step process: the mice were first rendered unconscious with inhalation of methoxyflurane (gas, 0.5ml) prior to cervical dislocation. Following euthanasia, the brain was rapidly dissected and placed into ice-cold cutting solution comprising and (mM) 206 sucrose, 2.5 KCl, 1.25 NaH_2_PO_4_, 25 NaHCO_3_, 25 glucose, 3 MgCl_2_, 1 CaCl_2_ and bubbled with carbogen (5% CO_2_/95% O_2_). Slices were kept at 37°C in a holding chamber containing artificial CSF (artificial cerebrospinal fluid-ACSF) comprising (mM) 125 NaCl, 3 KCl, 2 CaCl_2_, 1 MgCl_2_, 25 NaHCO_3_, 1.25 NaH_2_PO_4_ and 25 glucose bubbled with carbogen (5% CO_2_/95% O_2_). After incubation at 37°C for 1 hour, slices were held at room temperature until required.

### Electrophysiology

Slices were continuously perfused with ACSF (∼1.1ml/minutes) and maintained at 35 ± 2 °C (Warner Instruments TC-324B). For whole-cell current clamp recordings, 4-7 MΩ borosilicate glass patch electrodes (Harvard apparatus GC150F-15, 1.5 mm outer dimeter × 0.86 inner diameter) were filled with an internal solution comprising (mM) 135 potassium gluconate, 10 HEPES, 7 NaCl, 2 Na_2_ATP, 0.3 Na_3_GTP and 2 MgCl_2_.

Slices were visualized at 60× magnification under bright field and infrared differential interference contrast video microscopy (Olympus BX51-WI equipped an Oxford Instruments Andor Zyla 4.2 camera). Somatic recordings were made using a Sutter Double IPA Integrated Patch Amplifier (Sutter Instruments) under the control of SutterPatch (Sutter Instrument), with data acquired at a sampling rate of 50 kHz and low pass filtered at 10 kHz.

Whole cell current clamp recordings with no holding current were conducted on layer 2/3 and layer 5 pyramidal neurons. When possible, pyramidal neurons expressing eGFP were recorded from first (illuminated with CoolLED pE-300 ultra). To quantify AP properties (AP threshold, AP half-width, AP amplitude), single APs were evoked with a 2ms long depolarising current that increased in steps of +25pA until a single AP was induced.

To investigate evoked spike firing properties (I-F curves), APs were evoked with a current step protocol of 1s current steps ranging from -100 to + 500 pA (increasing in 25pA current steps with a 5s inter-step interval). This protocol was repeated for a second time after a 30s delay. Series resistance was checked prior to each current clamp recording. Neurons were excluded from analysis if the series resistance changed by >20% of the baseline value and/or exceeded 30 MΩ.

### Electrophysiological data analysis

Resting membrane potential (RMP) was the membrane potential at the start of the whole cell recording. Two single AP traces were analysed to get the average AP threshold (dV/dt of 10V/s), AP amplitude (relative to AP threshold), and AP half-width. Two I-F curve recordings were analysed to determine the average rheobase, input resistance, maximum spike firing frequency, and evoked spike firing frequency for each positive current step. For evoked spike firing frequency, the total number of AP’s fired in response to the 1s depolarising current steps was quantified. Representative AP and single AP traces are presented in Fig 1. D and E.

### Statistical analysis

We performed two-group estimation statistics to determine the mean differences in AIS structural and electrophysiological properties between stroke and sham-controlled mice. Cumming plots are presented detailing the mean differences between the two groups. Mean data is presented as mean ± 1 standard deviation, while mean differences are presented as mean difference ± 95% confidence intervals. Estimation statistical analysis and generation of figures were obtained using resources described by Ho et al. (2019).

A one-way Bayesian ANOVA was conducted to determine the amount of evidence for AIS structural and/or functional plasticity following stroke, with group (stroke vs no stroke), cortical layer (layer 2/3 vs layer 5), cortical hemisphere (peri-infarct zone of the ipsilesional hemisphere vs contralesional hemisphere), and sex (male vs female) as fixed factors. Bayesian statistics allows for a more informative conclusion as it provides continuous evidence for or against an effect (in the case of our study, in favour for the alternative hypothesis), leading to more reproducible science ^22^.

A custom prior value of 0.5 was used in our analysis. Bayes Factors (BF) are reported according to previously reported cut-offs that suggest evidence in favour of the alternative hypothesis: BF_10_ ≤1 = no evidence, 1 < BF_10_ ≤ 3= anecdotal evidence, 3 < BF _10_ ≤10= moderate evidence, 10 < BF _10_ ≤100= strong evidence, BF_10_ <100= extreme evidence. Where there was evidence for a main effect, a Bayesian post-hoc test based on pairwise comparisons using Bayesian t-tests was performed. Given the recency of Bayesian statistics compared to frequentist statistics, we also conducted one-way frequentist ANOVA to support our analysis ^22^. Where there was evidence for main effects and their interactions, post-hoc testing was performed using Tukey’s multiple comparison test and p values ≤ 0.05 were considered statistically significant. All Bayesian and Frequentist ANOVA results are presented in Supplementary Table 3.

Bayesian and frequentist statistical analysis were conducted using JASP (version 0.18.3). Normality was verified with visual inspection of Q-Q plots and homogeneity of variance tested with Levene’s test. Rheobase data was log transformed prior to analysis to conform with statistical assumptions of normality.

Due to an uneven sample size in the AIS structural analysis, sex was not included as a fixed factor for AIS distance from soma and the data was collapsed to assess changes between experimental group, cortical hemisphere, and cortical layer only.

## Results

### Pyramidal neuron AIS structure remains unaltered in the sub-acute phase post stroke

To determine if maladaptive AIS plasticity is present in the sub-acute phase post stroke, we first compared AIS structure (length and distance from soma) between stroke and sham animals. A Bayesian ANOVA indicated no evidence for an effect of group (BF_10_=0.437, BF_incl_=0.299) on AIS length in the sub-acute phase post stroke with a -0.015 µm [95.0%CI - 2.47, 2.23] unpaired mean difference between stroke and sham mice (Fig 2. A). Additionally, there was no evidence for an effect of layer (BF_10_=0.388, BF_incl_=0.29; layer 2/3 mean length: 21.99 ± 2.86 µm (n=21 cells); layer 5 mean length:21.33 ± 2.71 µm (n=21 cells); unpaired mean difference:-0.657 [95.0%CI -2.3, 0.991]), or sex (BF_10_=0.390, BF_incl_=0.111; male mean length:21.95 ± 2.66 µm (n=24 cells); female mean length:21.27 ± 2.95 µm (n=18 cells); unpaired mean difference:0.68 [95.0%CI -0.981, 2.42]) on AIS length. There was anecdotal evidence for an effect of hemisphere on AIS length irrespective of the group condition (BF_10_=1.935, BF_incl_=1.59; contralesional mean length:22.4 ± 2.85 µm (n=25 cells); ipsilesional mean length:20.57 ± 2.33 µm (n=17 cells); unpaired mean difference: -1.82 µm [95.0%CI - 3.36, -0.248]; Fig 2. B).

**Figure 2.**
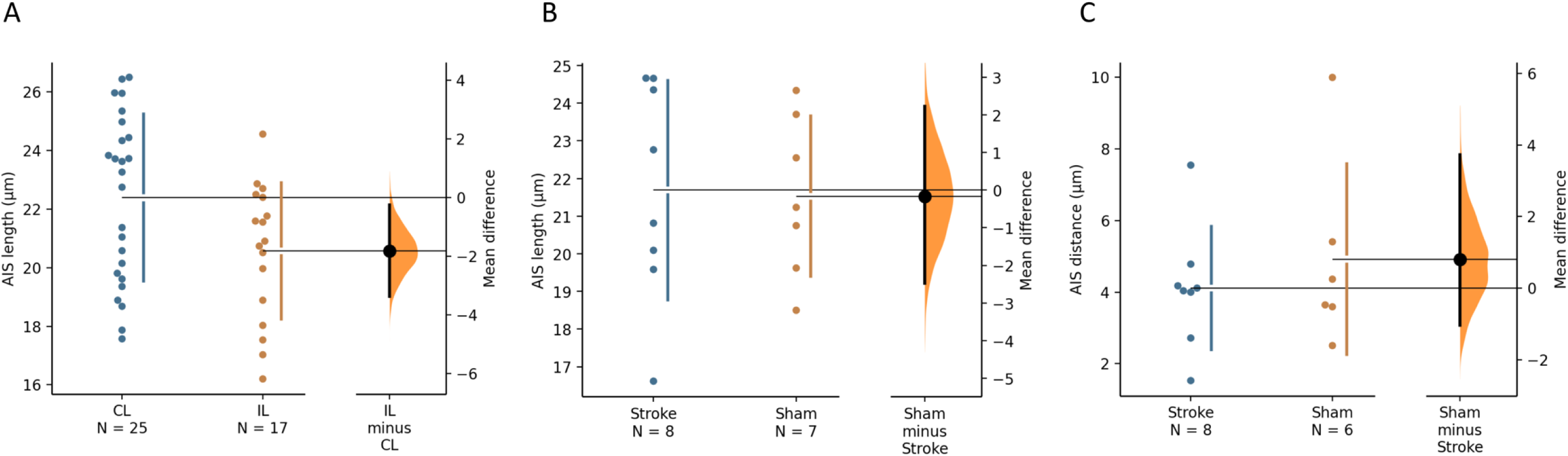
Cumming estimation plots of AIS length and AIS distance from the soma at 28-day post stroke. There was no evidence of a change in AIS length (A) or AIS distance (B) between stroke and sham controlled mice at the sub-acute phase of post stroke. There was anecdotal evidence for shorter AIS lengths in the peri-infarct zone of the ipsilesional (IL) hemisphere relative to the contralesional (CL) hemisphere (C).

Similar to our findings on AIS length, AIS distance from the soma appeared consistent between stroke (4.14 ± 1.73 µm, n=6 mice) and sham mice (4.92 ± 2.67 µm, n=6 mice), with an unpaired mean difference of 0.781 µm [95.0%CI -0.952, 3.81] (Fig 2. C). A Bayesian ANOVA revealed no evidence for an effect of group (BF_10_=0.524, BF_incl_=0.234), hemisphere (BF_10_=0.55, BF_incl_=0.262; contralesional mean distance: 3.88 ± 1.95 µm (n=20 cells); ipsilesional mean distance:4.62 ± 1.63 µm (n=13 cells); unpaired mean difference:0.598 [95.0%CI -0.685, 1.81]), or cortical layer (BF_10_=0.379, BF_incl_=0.197; layer 2/3 mean distance: 3.98 ± 2.0 µm (n=16 cells); layer 5 mean distance:4.35 ± 1.71 µm (n=17 cells); unpaired mean difference:0.348 [95.0%CI -0.967, 1.62]) on AIS distance. Taken together, these findings show that AIS structure remains intact in the sub-acute phase post stroke.

### Pyramidal neuron AP threshold, spike properties, and RMP are unaltered in the sub-acute phase post stroke

Despite observing no alterations in AIS structure, we still sought to characterise AIS function and intrinsic excitability properties in the sub-acute phase post stroke. In line with our findings AIS structural measures, our analysis revealed no evidence for an effect of group on AP threshold (stroke mean: -44.4 ± 4.28 mV; sham mean: -44.1 ± 4.43 mV; unpaired mean difference: -0.028 mV [95.0%CI -1.57, 1.16]; BF_10_=0.215, BF_incl_=0.066, Fig 3. A), AP half-width (stroke mean:0.577 ± 0.169 ms; sham mean:0.611 ± 0.204 ms; unpaired mean difference:0.025 ms [95.0%CI -0.046, 0.101]; BF_10_=0.304, BF_incl_=0.057, Fig 3. B), AP amplitude (stroke mean:90.7 ± 8.9 mV; sham mean: 91.9 ± 7.2 mV; unpaired mean difference:1.68 mV [95.0%CI -1.33, 4.72]; BF_10_=0.265, BF_incl_=0.071, Fig 3. C), or rheobase (stroke mean:185.1 ± 115.3 pA; sham mean:159.0 ± 83.7 pA; unpaired mean difference: -24.4 pA [95.0%CI -64.5, 11.9]: BF_10_=0.477, BF_incl_=0.074, Fig 3. D). Analysis of other intrinsic membrane measures showed no evidence for an effect of group on RMP (stroke mean: -67.7 ± 5.36 mV; sham mean: -68.6 ± 57 mV; unpaired mean difference: -0.848 mV [95.0%CI -2.78, 1.03]; BF_10_=0.284, BF_incl_=0.059, Fig 3. E), or maximum spike firing frequency (stroke mean:31.6 ± 15.1 Hz; sham mean:37.0 ± 15.9 Hz; unpaired mean difference:6.24 Hz [95.0%CI 0.393, 12.0]; BF_10_=0.845, BF_incl_=0.264, Fig 3. F)

**Figure 3:**
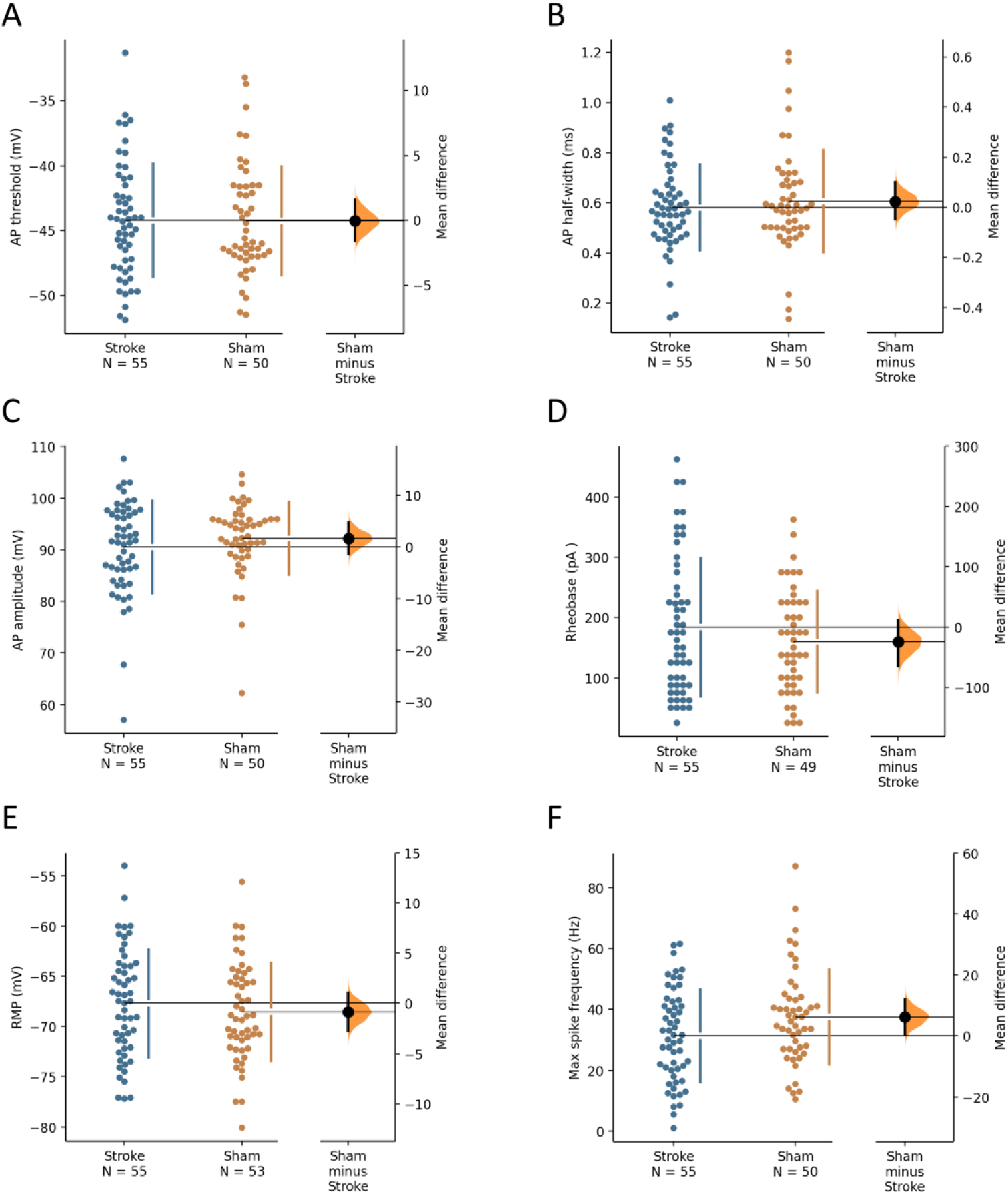
Cumming estimation plots of intrinsic neuronal properties at 28-days post stroke. No changes were observed between stroke and sham controlled mice in AP threshold (A), AP half-width (B), AP amplitude (C), rheobase (D), RMP (E) and maximum spike frequency (F). The number of cells analysed in each group is presented under the group names.

While we found no evidence for a group-dependent change in AIS firing properties, we did find a sex-dependent effect (i.e. in both stroke and sham) on AP threshold and AP amplitude. Male mice had a slightly more depolarised AP threshold compared to female mice (male mean: -44.95 ± 3.86 mV; female mean: -43.27 ± 4.61 mV, n=60 cells; unpaired mean difference of 1.85 mV [95.0%CI 0.215, 3.57]; Supplementary Fig 1. A), and a reduced AP amplitude compared to female mice (male mean:89.58 + 8.51 mV; female mean:92.63 + 7.62 mV; unpaired mean difference of -3.52 mV [95.0%CI -6.87, -0.637]; supplementary Fig 1. B). Our Bayesian ANOVA indicated anecdotal evidence for an effect of sex on AP threshold (BF_10_=1.286, BF_incl_=0.264) and AP amplitude (BF_10_=1.075, BF_incl_=0.361. Moreover, there was anecdotal evidence for an effect of sex on RMP (BF_10_=1.791, BF_incl_=0.369). Female mice had a slightly more hyperpolarised RMP relative to male mice (female mean: -69.1 ± 4.34 mV; male mean: -66.9 ± 5.82 mV; unpaired mean difference 2.2 mV [95.0%CI 0.165, 4.12]; supplementary Fig 1. C).

Our analysis also revealed layer-dependent changes in AP half-width and maximum spike firing frequency irrespective of group. Our Bayesian analysis revealed moderate evidence for an effect of layer on AP half-width (BF_10_=3.469, BF_incl_=0.685) and maximum spike firing frequency (BF_10_=7.771, BF_incl_=2.069), with posterior odds of 3.469 and 7.771, respectively. Further inspection showed that layer 2/3 pyramidal neurons displayed a wider AP half-width on average (layer 2/3 mean:0.644 + 0.21 ms; layer 5 mean:0.553 + 0.02 ms; unpaired mean difference: -0.091ms [95.0%CI -0.164, -0.021; supplementary Fig 1. D) and attenuated maximum spike firing frequencies (layer 2/3 mean:29.4 ± 14.1 Hz; layer 5 mean:37.9 ± 15.8 Hz; unpaired mean difference:8.300 Hz [95.0%CI 2.67, 14.0]; supplementary Fig 1. E) relative to layer 5 pyramidal neurons.

### Stroke mice show decreased input resistance in the sub-acute phase of stroke

While we found no stroke-induced changes in AP threshold, spike properties, or RMP in pyramidal neurons, we did find a difference in input resistance in the sub-acute stage post stroke. Stroke mice had a mean input resistance of 63.07 ± 28.96 MΩ (n=55 cells) compared to 77.9 ± 36.04 MΩ in sham-controlled mice (n=51 cells), with an unpaired mean difference of 14.8 MΩ [(95.0%CI 2.59, 27.4; Fig.4]. Our Bayesian ANOVA indicated anecdotal evidence for an effect of group on input resistance (BF_10_=2.311, BF_incl_=0.364, Fig 4). There was no evidence for an effect of sex, hemisphere, or layer on input resistance. These results suggest that stroke may decrease input resistance in the sub-acute stage post stroke.

**Figure 4:**
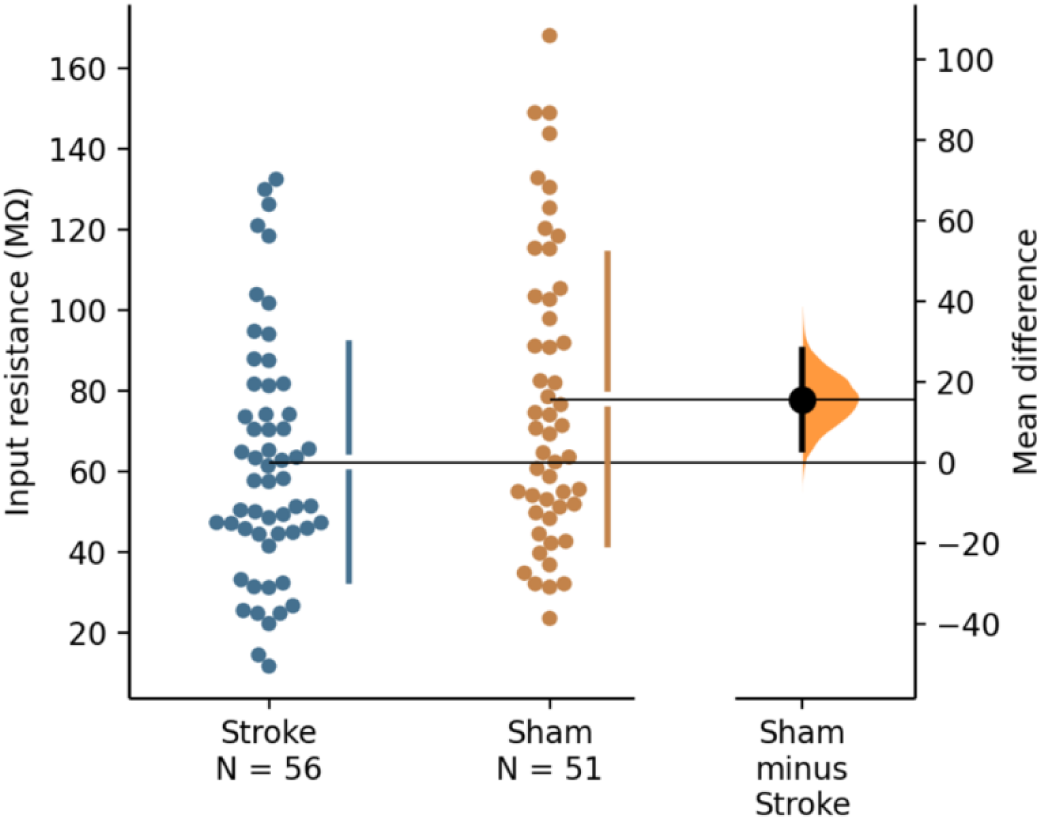
Cumming estimation plot of input resistance at 28-days post stroke. There was anecdotal evidence for an effect of stroke on input resistance with stroke mice showing reduced input resistance relative to sham mice in the sub-acute phase post stroke. The number of cells analysed in each group is presented under the group names.

**Figure 5.**
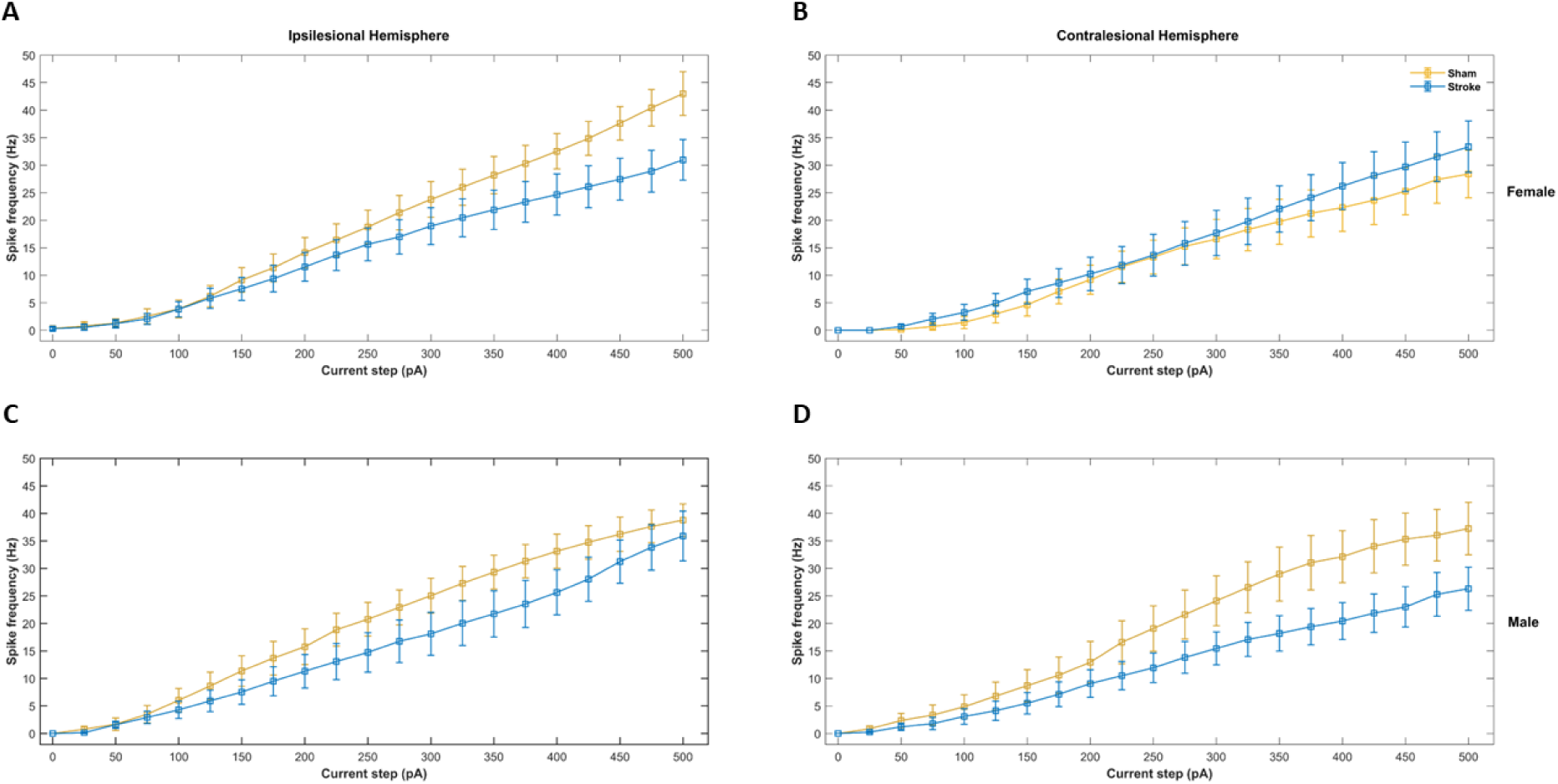
IF curves of evoked spike firing frequency. Stroke reduced evoked spike firing in the ipsilesional and contralesional hemispheres in male mice (C-D) and in the ipsilesional hemisphere of female mice (A) relative to sham controlled mice.

### Pyramidal neuron evoked spike firing frequency is attenuated in the sub-acute phase post stroke

As a broad measure of intrinsic excitability, we quantified the evoked spike frequency in response to increasing steps of depolarising current (Fig 4: I-F curves – see Supplementary Table 7 for summary of descriptive statistics). Analysis of the I-F curves showed extreme evidence for an effect of group, sex, hemisphere, cortical layer, and current step on the evoked spike frequency (BF_10_=∞ and BF_incl_>600 for all factors) (see Supplementary Table 8 for summary of estimates). In addition, there was evidence for the interactions group*sex*hemisphere (BF_incl_=16.50) and group*sex*layer (BF_incl_=463.56). As there was strong evidence for the main effects and some of their interactions, which was mirrored by our frequentist ANOVA analysis (See supplementary Table 9 for summary of ANOVA), we interpreted these effects with the aid of frequentist Tukey post-hoc testing of the interactions rather than post-hoc testing of the main effects. Post-hoc testing of the group*sex*hemisphere interaction showed stroke reduced the evoked spike firing frequency compared to sham in the peri-infarct zone of the ipsilesional hemisphere in females (mean difference=-4.10, t=4.616, p<0.001) and males (mean difference=-4.64, t=4.43, p<0.001), as well as in the contralesional hemisphere, but only in males (mean difference=-6.768, t=7.14, p<0.001). In the contralesional hemisphere of sham mice, evoked spike firing frequency was reduced in females compared to males (mean difference=-6.24, t=6.543, p<0.001). Post-hoc testing of the group*sex*cortical layer interaction showed that the evoked spike firing in layer 2/3 of sham mice was reduced in females compared to males (mean difference=-6.44, t=6.149, p<0.001).

## Discussion

Here we characterised whether maladaptive AIS and intrinsic plasticity occurs in the sub-acute phase post stroke how this varied between the ipsilesional and conrtralesional hemisphere, and sex. Surprisingly we found that normal AIS structure and function is maintained in the sub-acute phase post stroke in both male and female mice. We did however find decreased input resistance and pyramidal neuron excitability in the peri-infract hemisphere in both sexes. In addition, reduced pyramidal neuron excitability extended to the contralesional hemisphere, but only in males.

In the acute phase post stroke (0 – 2 weeks), it is known that the AIS of surviving neurons adjacent to the peri-infarct zone undergo rapid proteolysis over the first 72 hours post-injury ^18^ and shortening of ∼3µm in length across all cortical layers by 2 weeks post stroke ^19^. Our study extends these findings by characterising both AIS structure and function in the sub-acute phase post stroke when neural plasticity is in a heightened state and rehabilitation typically begins. In contrast to the maladaptive structural changes seen in the acute phase, we found that AIS length and distance from soma were not different from sham in layer 2/3 or 5. This was further supported by our electrophysiological measures of AIS function which showed no differences in AP threshold or rheobase between stroke and sham. This suggests that ischemic stroke leads to maladaptive AIS structural plasticity in the acute phase that returns to normal in the sub-acute phase. Interestingly our results reflect a similar pattern seen with post stroke axonal sprouting which occurs over the first 2 weeks post stroke but subsides by day 28 ^23^. Therefore, we speculate that structural axonal plasticity, both beneficial and maladaptive, is recruited during the acute phase of stroke but does not continue into the sub-acute phase. From a rehabilitation point of view, this quite promising as it suggests maladaptive AIS plasticity is temporary. However, while our results show restoration of normal AIS structure and function, further research is needed to determine whether the AIS can still undergo normal activity-dependent plasticity. This will be important as normal AIS plasticity is required to keep neuronal activity within homeostasis in adulthood^24^. Furthermore, the efficacy of stroke treatments that aim to promote functional recovery by manipulating neural activity, such as non-invasive brain stimulation^25–27^ could be reduced if the capacity for AIS plasticity is impaired.

Despite normal AIS structure and function, we found further evidence of reduced pyramidal neuron excitability in the peri-infract area and contralesional hemisphere in the sub-acute phase post stroke ^28–31^. While reduced excitability, particularly in the peri-infarct zone, is mostly attributed to synaptic mechanisms and increased GABAergic inhibition ^11,28^, the reduced evoked spike firing frequency in our study is independent of synaptic changes and due to intrinsic membrane properties. The stroke-induced change to intrinsic excitability appears to be specific as the only intrinsic property that showed evidence of a change following stroke was input resistance. Interestingly a reduction in input resistance has previously been shown to occur in sub-cortical neurons, also during the sub-acute phase of stroke ^15^. Given that our results are from cortical pyramidal neurons, we suggest that reduced input resistance could be a general intrinsic property affected by ischemic stroke.

A major finding of our study was the sex-dependent reduction in pyramidal neuron excitability following stroke, with the changes confined to the peri-infarct hemisphere in female mice, whereas male mice showed changes in both the peri-infract and contralesional hemispheres. While previous rodent studies have shown sex-dependent differences in the extent of tissue damage ^32^ and microglial activation ^33^, we show that sex differences also occur to neuronal function. Similar to previous reports, we find that stroke had a greater impact on male mice, providing further support for a beneficial role of oestrogen ^34,35^. However, it is important to note that in our study, the greater impact relates to the distance/number of cortical circuits affected by stroke and not the effect size as the reduction in evoked spike firing frequency in the peri-infract hemisphere following stroke was almost identical in males and females. The underlying mechanism of this sex difference remains unclear as almost all the intrinsic properties that showed anecdotal evidence of a difference between sex, irrespective of the injury condition (RMP, AP threshold and AP amplitude), were not the same properties altered by stroke. While we found reduced spike firing frequencies in layer 2/3 of sham female mice compared to sham male mice, this occurred across both hemispheres and therefore does not explain why changes to the contralesional hemisphere is unique to male mice. Nevertheless, our study provides important insight into the sex-dependent differences of stroke pathology at the cellular level and emphasises the need to develop personalised treatments for ischemic stroke patients.

A limitation of our study is the inability to correlate our structural and electrophysiological measures with behaviour (e.g., forelimb function). For our study, we used a photothrombotic stroke protocol known to induce motor function deficits from the first day post stroke in the age and species of mouse used ^20^, even at lower Rose Bengal and illumination times ^36^. Therefore, it would be useful for future studies to correlate AIS changes to behavioural changes, in both the acute and sub-acute phases post stroke to determine a potential relationship between maladaptive AIS plasticity and the extent of neurological impairment. Another limitation is the unequal sample size in some of our outcome measures. In particular, the unequal sample size of our electrophysiology due to the exclusion of a small number of inhibitory cells that were also transfected under the CaMKII promoter ^37–39^. While the removal of cells was not specific to particular groups, this may have impacted our ability to determine differences in specific cortical layers of each hemisphere (e.g., layer 2/3 in the contralesional hemisphere of males). Unfortunately, the number of inhibitory cells recorded in each group only allowed us to analyse a small number of inhibitory neurons at the group level (i.e. stroke vs sham). Our analysis of this small sample shows no change to AIS or other intrinsic excitability measures in the sub-acute phase post stroke but confirmation with larger sample sizes is needed, especially for hemisphere, cortical layer, and sex dependent analyses.

In conclusion, our results show that despite normal structure and function of the AIS, neural excitability is reduced in pyramidal neurons across both sexes through other intrinsic membrane properties. Our findings have important implications for the treatment of ischemic stroke and suggest that interventions targeting AIS dysfunction should be applied in the acute phase rather than the sub-acute phase. Furthermore, that changes to intrinsic excitability, in part, underlies the reduced neuronal excitability observed in the sub-acute stage post stroke, which requires different management between males and females.

## Acknowledgments

We would like to acknowledge Liz Jaeschke-Angi for her technical assistance.

## Supplementary Materials

Supplementary results

Tables S1-S9

Figures S1

## Lesion volume analysis

**Table 1.** Stroke lesion volume. * Not included in structural analysis due to insufficient data.

| Animal ID | Sex | Lesion size (mm<sup>3</sup>) |
| --- | --- | --- |
| 010* | Female | 1.34 |
| 016 | Male | 0.68 |
| 021* | Female | 0.72 |
| 025 | Male | 1.05 |
| 027 | Male | 1.15 |
| 047 | Male | 0.85 |

The stroke lesion volume was quantified for 7 mice (Fig 1. A) and revealed substantial variability between mice (table 1).

## Electrophysiological cell counts

**Table 2.** Electrophysiological cell counts split by sex, hemisphere, and layer for stroke and sham groups. Where possible, we endeavoured to obtain a minimum of one cellular recording per layer (i.e layer 2/3 and layer 5) of each hemisphere (contra- and ipsilesional) for every animal (12 mice total, split evenly by sex).

| Sex | Hemisphere | Layer | Group | # Cells |
| --- | --- | --- | --- | --- |
| Female | Contralesional | 2/3 | Sham | 6 |
|  |  |  | Stroke | 6 |
|  |  | 5 | Sham | 7 |
|  |  |  | Stroke | 7 |
|  | Ipsilesional | 2/3 | Sham | 8 |
|  |  |  | Stroke | 7 |
|  |  | 5 | Sham | 11 |
|  |  |  | Stroke | 10 |
| Male | Contralesional | 2/3 | Sham | 5 |
|  |  |  | Stroke | 3 |
|  |  | 5 | Sham | 6 |
|  |  |  | Stroke | 9 |
|  | Ipsilesional | 2/3 | Sham | 6 |
|  |  |  | Stroke | 6 |
|  |  | 5 | Sham | 4 |
|  |  |  | Stroke | 7 |

## Preliminary data suggests inhibitory neuron intrinsic excitability remain unchanged in the sub-acute phase post stroke

AAV transfection with the CaMKII promoter leads to labelling in a small proportion of inhibitory neurons. As a result, we collected whole-cell electrophysiological data from inhibitory neurons when targeting GFP positive cells. Due to the small sample size, we restricted our preliminary analysis to the group level only (i.e. stroke vs sham – see Table 2 for descriptive statistics). Analysis with Bayesian ANOVAs found no evidence (BF<1) for a difference between sham and stroke for RMP, AP threshold, AP half-width, AP amplitude, rheobase, input resistance, or max spike firing frequency.

**Table 3.**
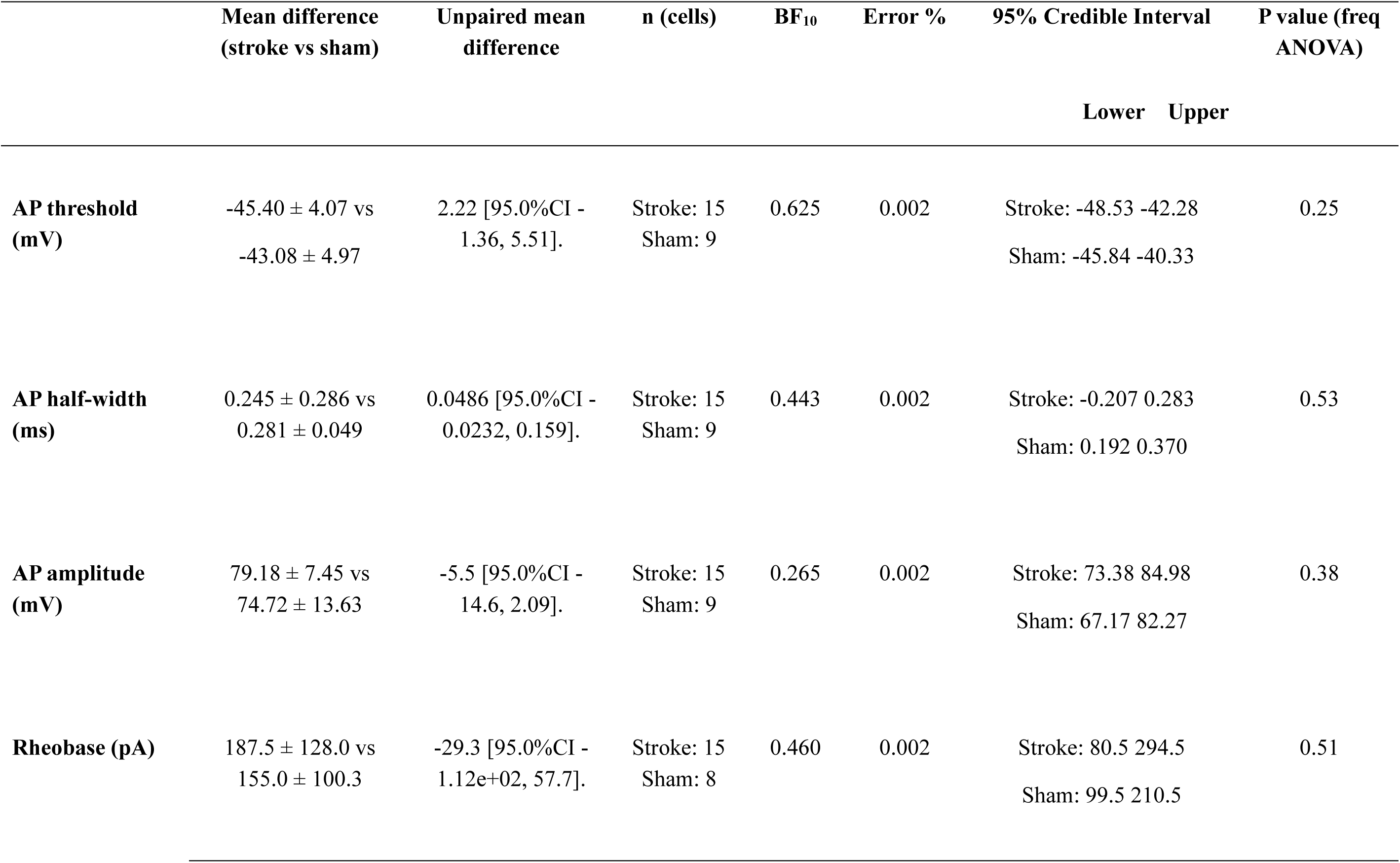
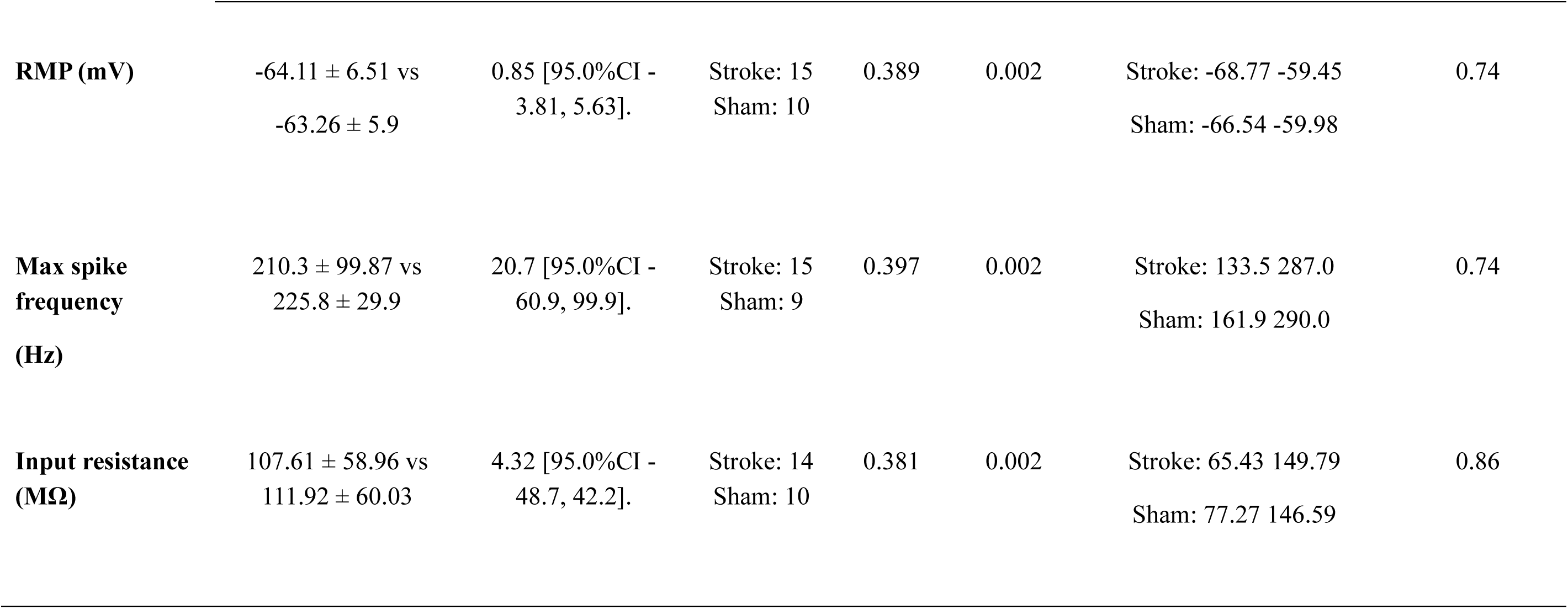
Summary of inhibitory neuron electrophysiological measures.

|  | Mean difference<br>(stroke vs sham) | Unpaired mean<br>difference | n (cells) | BF<sub>10</sub> | Error % | 95% Credible Interval | P value (freq<br>ANOVA) |
| --- | --- | --- | --- | --- | --- | --- | --- |
|  |  |  |  |  |  | Lower Upper |  |
| **AP threshold**<br>**(mV)** | -45.40 ± 4.07 vs<br>-43.08 ± 4.97 | 2.22 [95.0%CI -<br>1.36, 5.51]. | Stroke: 15<br>Sham: 9 | 0.625 | 0.002 | Stroke: -48.53 -42.28<br>Sham: -45.84 -40.33 | 0.25 |
| **AP half-width**<br>**(ms)** | 0.245 ± 0.286 vs<br>0.281 ± 0.049 | 0.0486 [95.0%CI -<br>0.0232, 0.159]. | Stroke: 15<br>Sham: 9 | 0.443 | 0.002 | Stroke: -0.207 0.283<br>Sham: 0.192 0.370 | 0.53 |
| **AP amplitude**<br>**(mV)** | 79.18 ± 7.45 vs<br>74.72 ± 13.63 | -5.5 [95.0%CI -<br>14.6, 2.09]. | Stroke: 15<br>Sham: 9 | 0.265 | 0.002 | Stroke: 73.38 84.98<br>Sham: 67.17 82.27 | 0.38 |
| **Rheobase (pA)** | 187.5 ± 128.0 vs<br>155.0 ± 100.3 | -29.3 [95.0%CI -<br>1.12e+02, 57.7]. | Stroke: 15<br>Sham: 8 | 0.460 | 0.002 | Stroke: 80.5 294.5<br>Sham: 99.5 210.5 | 0.51 |
| **RMP (mV)** | -64.11 ± 6.51 vs<br>-63.26 ± 5.9 | 0.85 [95.0%CI -<br>3.81, 5.63]. | Stroke: 15<br>Sham: 10 | 0.389 | 0.002 | Stroke: -68.77 -59.45<br>Sham: -66.54 -59.98 | 0.74 |
| **Max spike**<br>**frequency**<br>**(Hz)** | 210.3 ± 99.87 vs<br>225.8 ± 29.9 | 20.7 [95.0%CI -<br>60.9, 99.9]. | Stroke: 15<br>Sham: 9 | 0.397 | 0.002 | Stroke: 133.5 287.0<br>Sham: 161.9 290.0 | 0.74 |
| **Input resistance**<br>**(MΩ)** | 107.61 ± 58.96 vs<br>111.92 ± 60.03 | 4.32 [95.0%CI -<br>48.7, 42.2]. | Stroke: 14<br>Sham: 10 | 0.381 | 0.002 | Stroke: 65.43 149.79<br>Sham: 77.27 146.59 | 0.86 |

## Bayesian and Frequentist ANOVA results

**Table 4.**
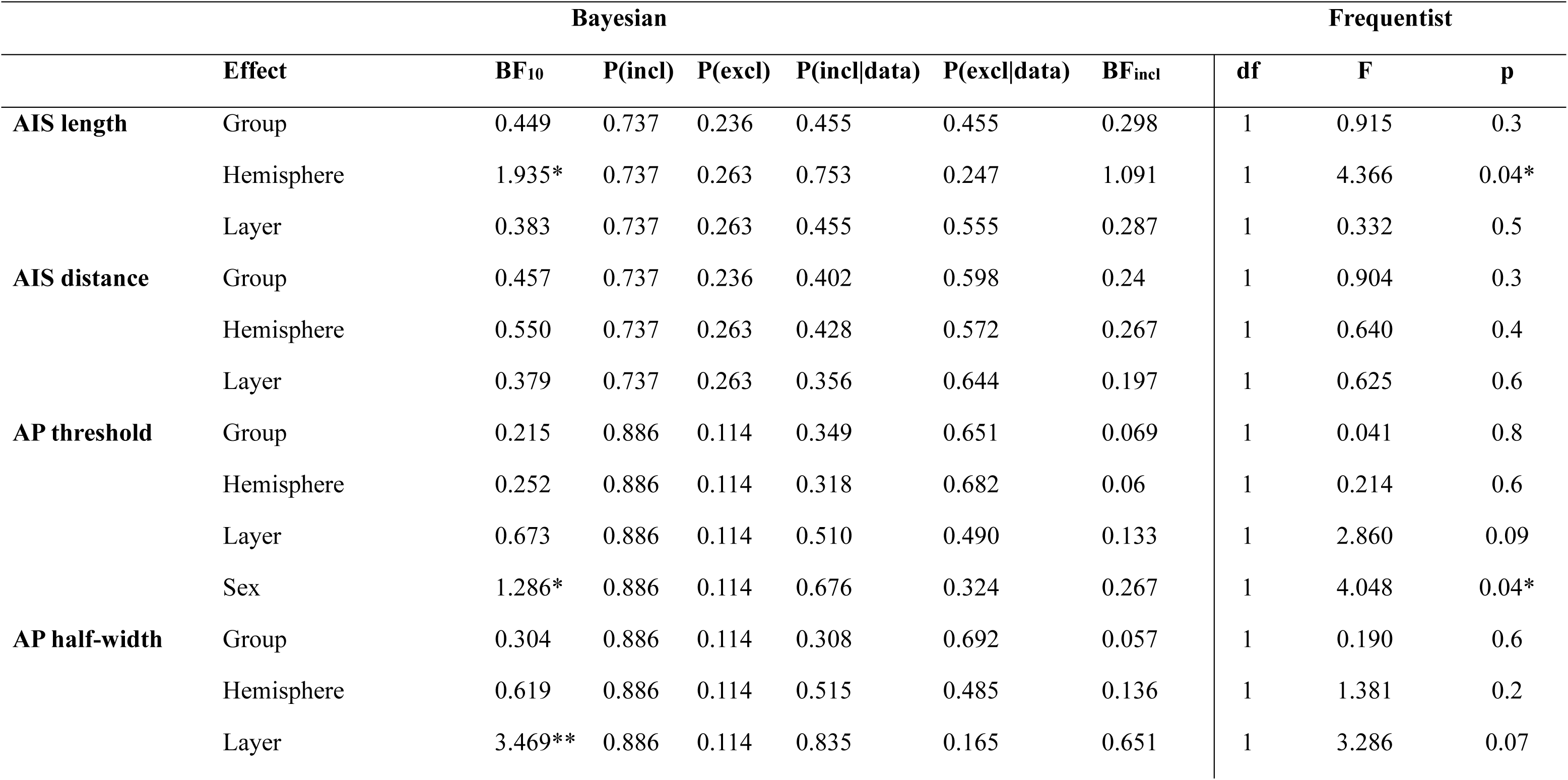
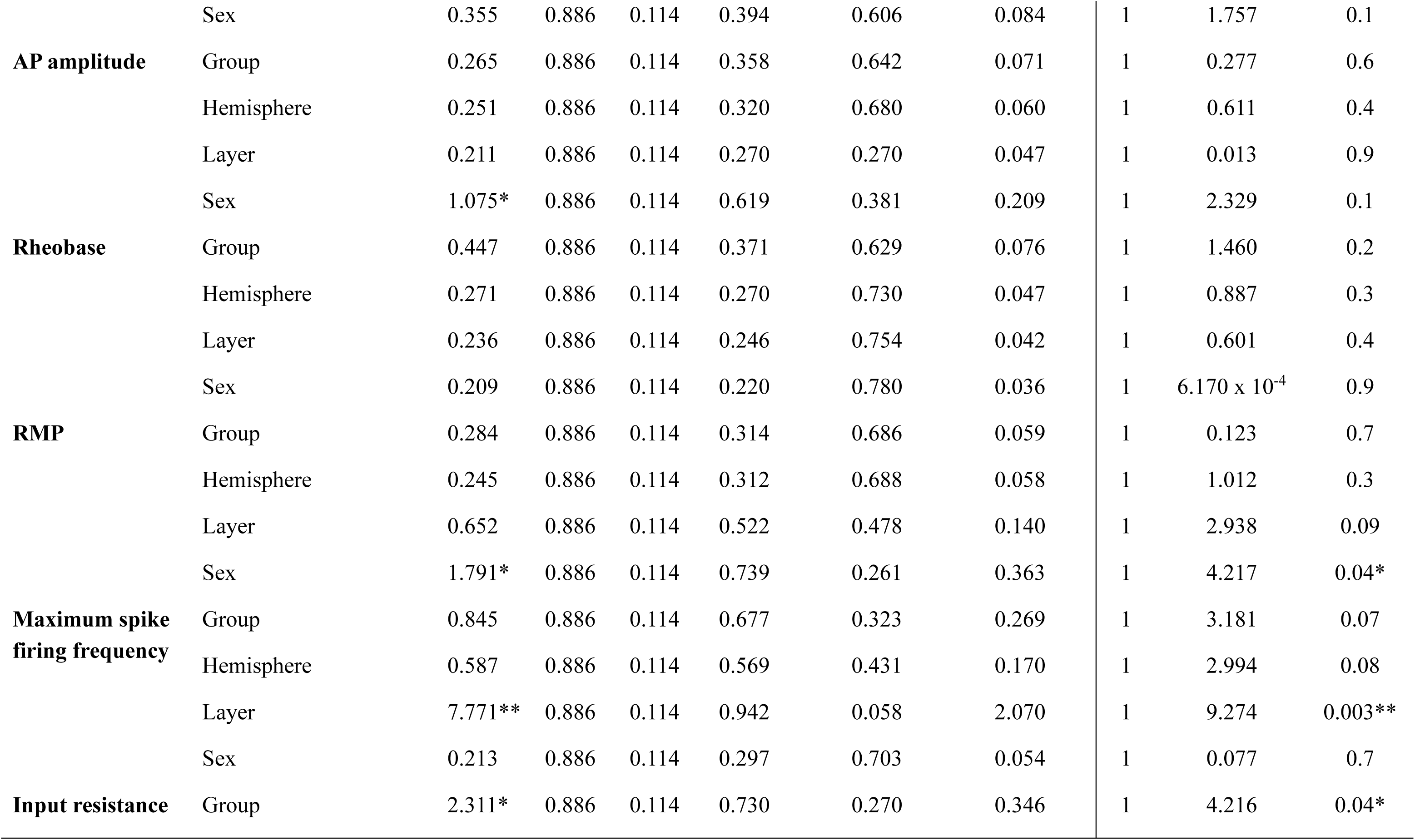
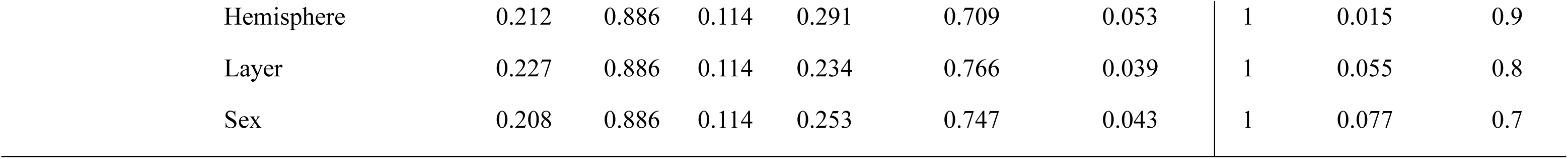
Bayesian and Frequentist ANOVA results for structural and electrophysiological changes in the sub-acute phase post stroke. For evoked spike firing frequencies, see table 7-8.

**Table 5.**
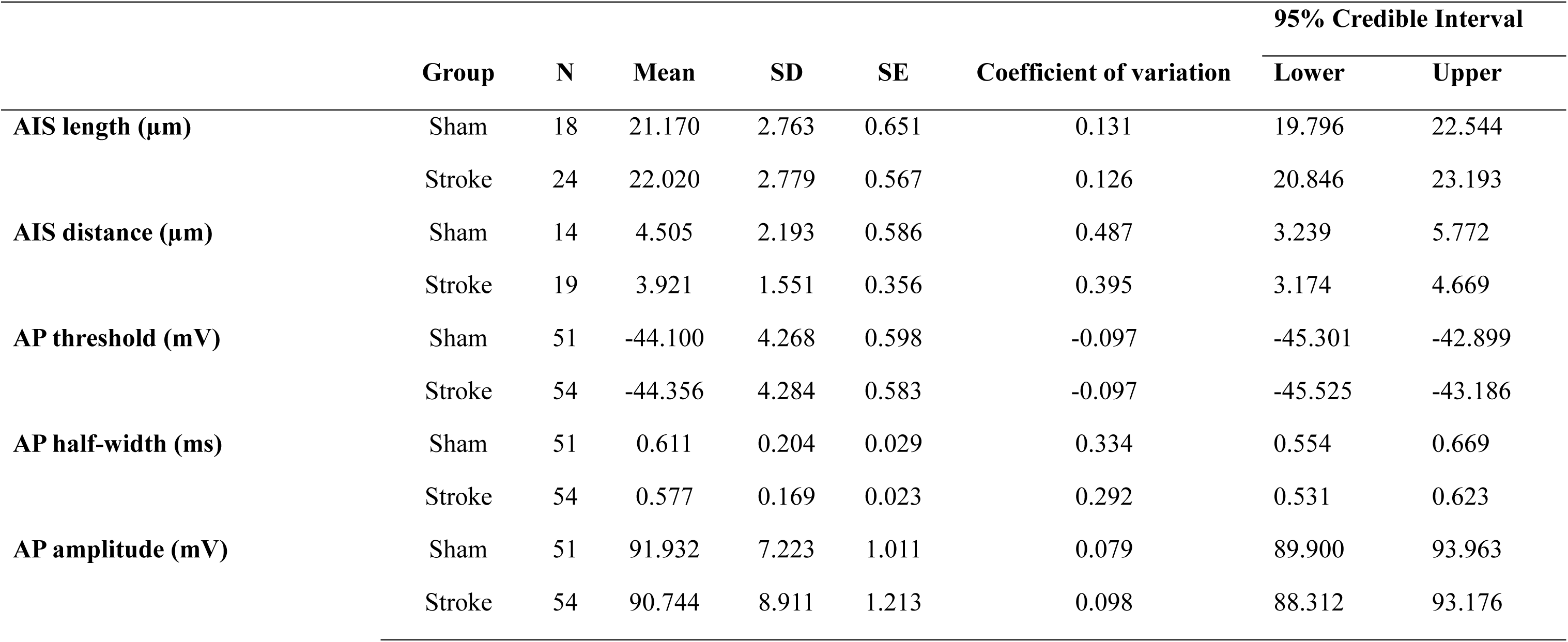
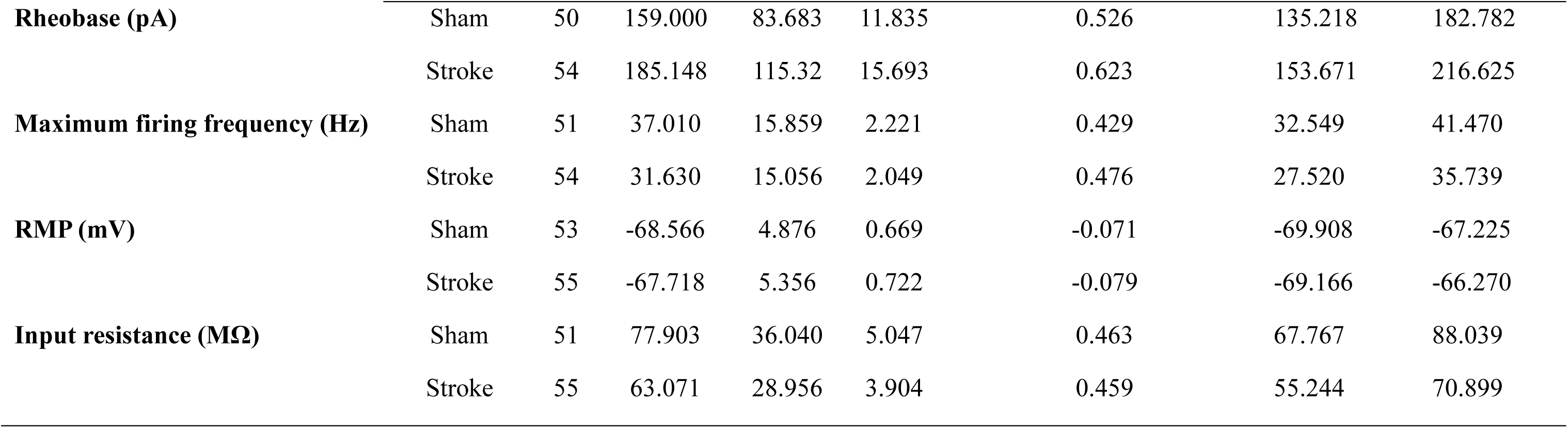
Descriptives table for all structural and electrophysiological changes observed at the group level in the subacute phase of post stroke. Mean differences are presented as the mean ± standard deviation.

**Table 6.**
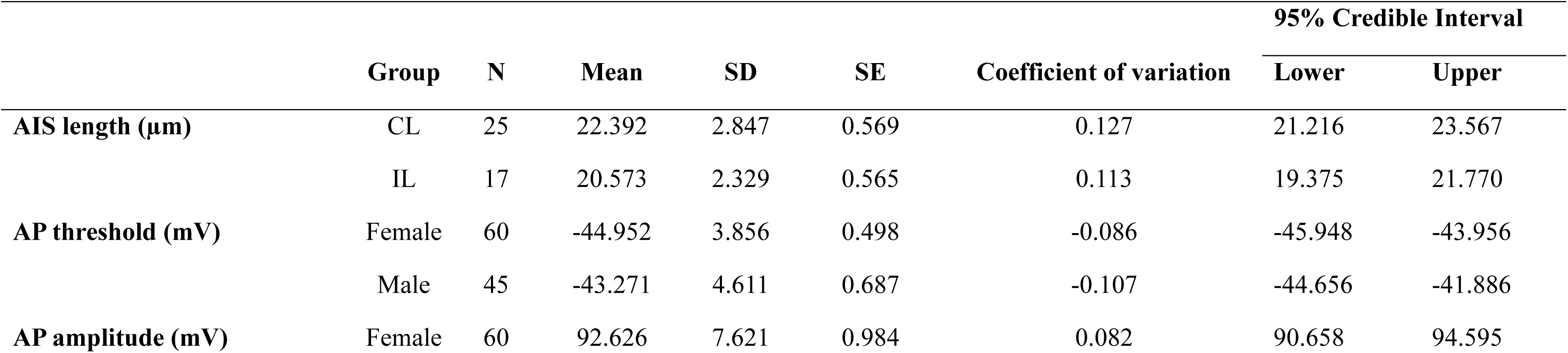
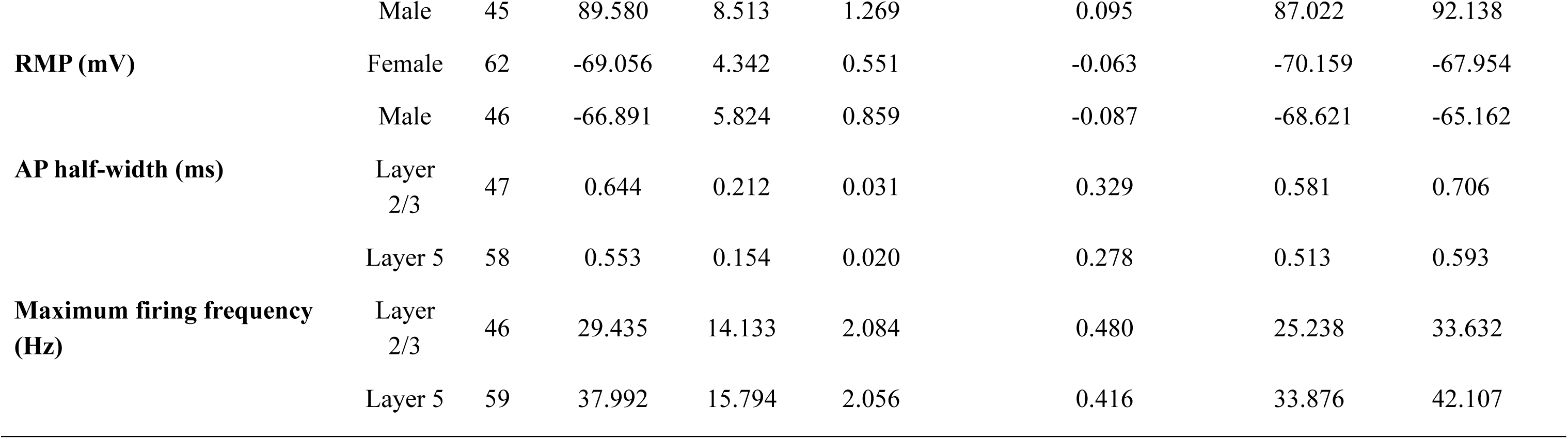
Descriptives table for structural and electrophysiological changes with evidence for an effect of hemisphere, layer or sex in the subacute phase of post stroke. Mean differences are presented as the mean ± standard deviation.

**Table 7.**
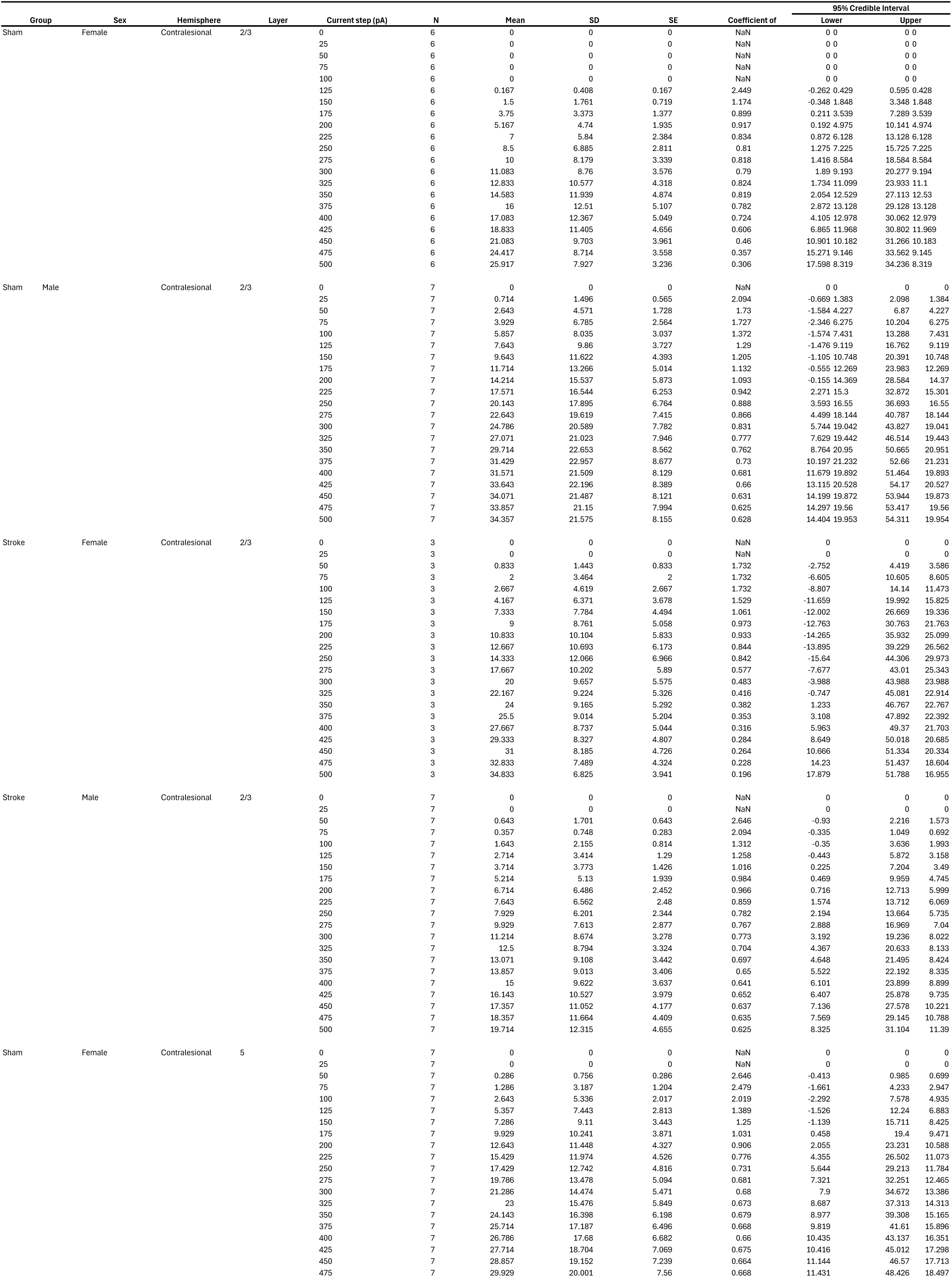
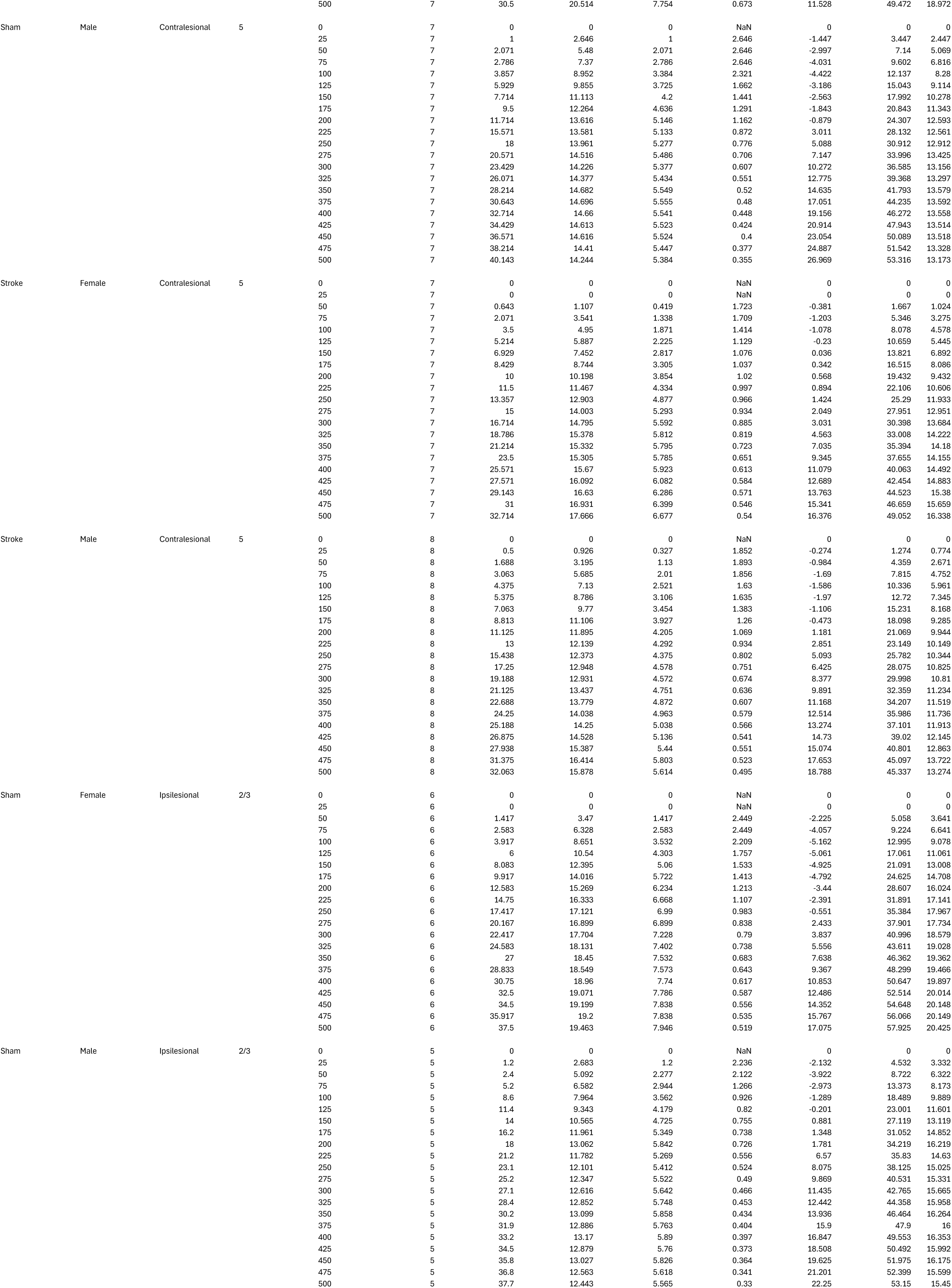
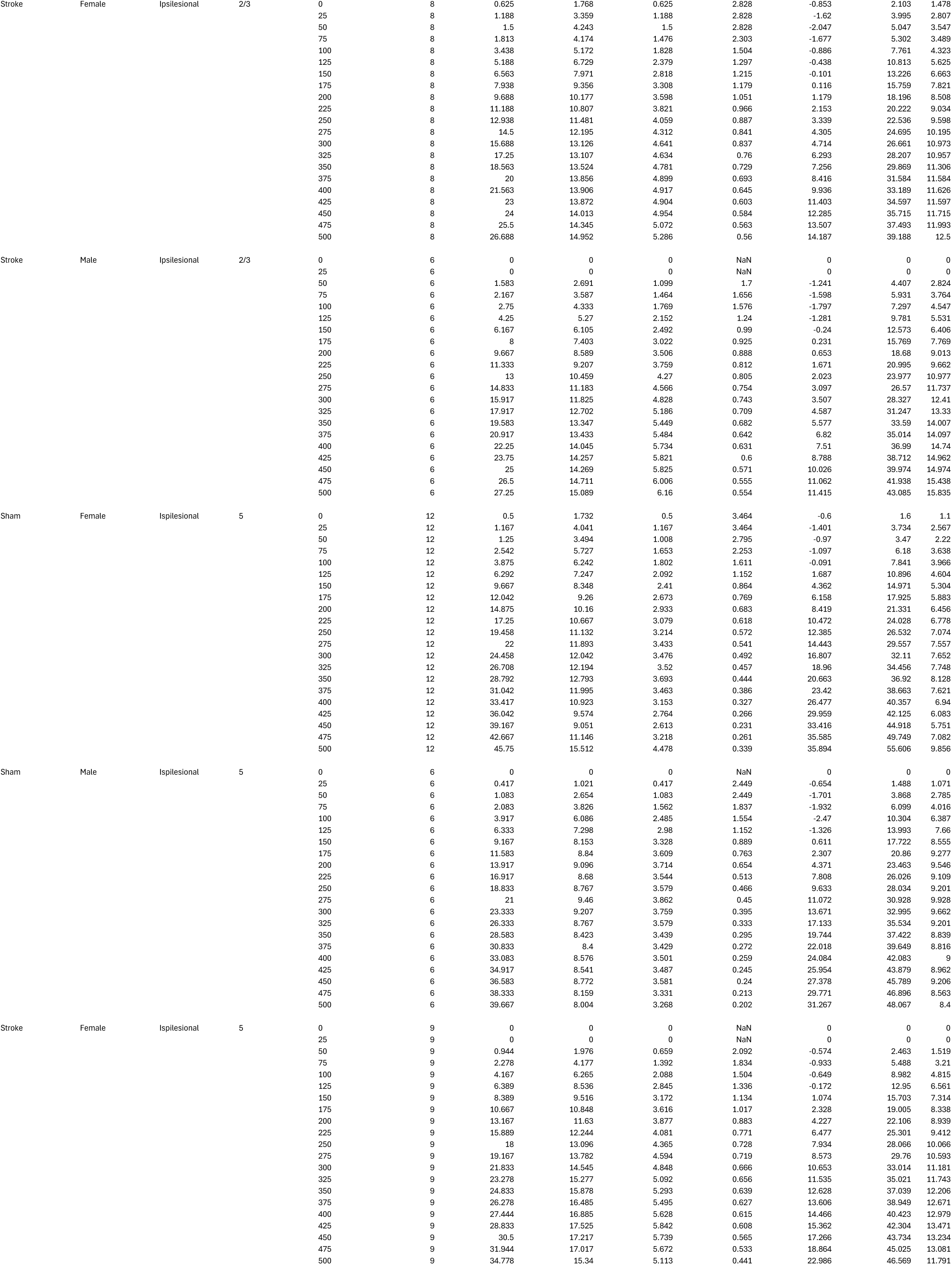
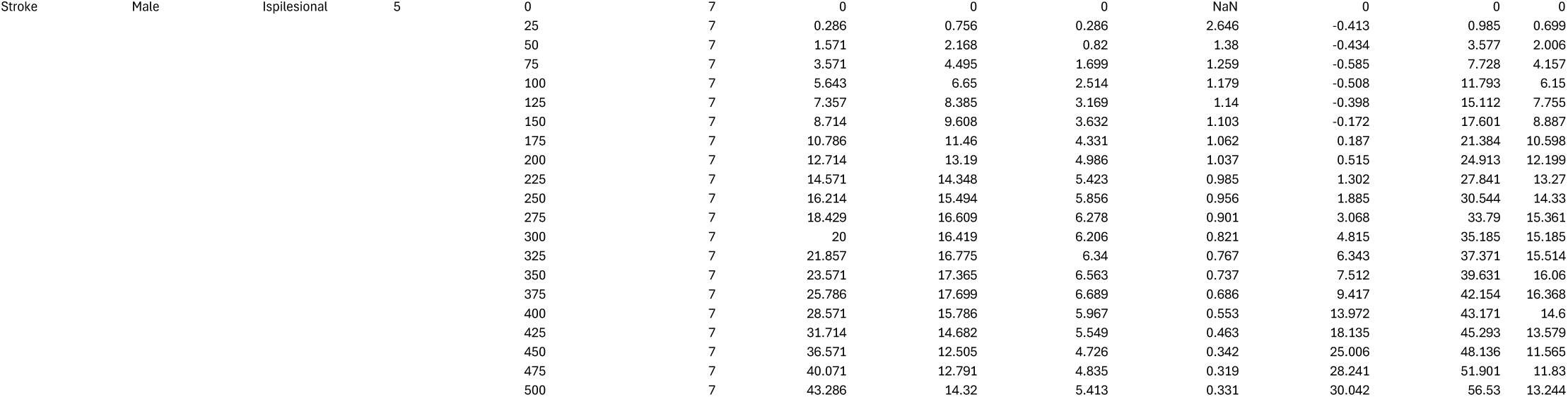
Descriptive statistics for IF curves.

**Supplementary Figure 1.**
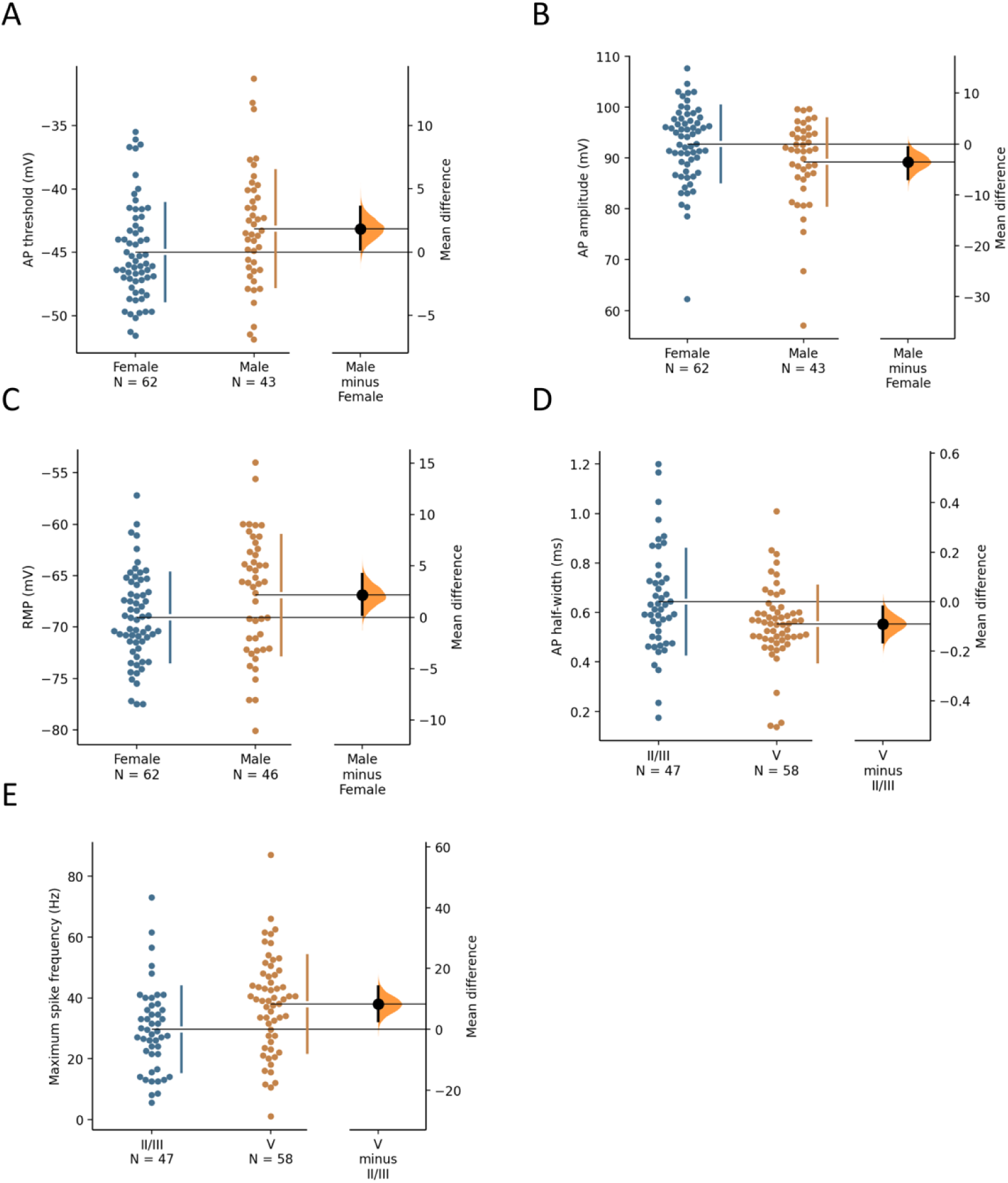
Cumming estimation plots of hemisphere, sex, and cortical layer-dependent changes in AIS and intrinsic neuronal properties. There was anecdotal evidence for a more hyperpolarised AP threshold (A) and increased AP amplitude (B) in female mice, while male mice had a more depolarised RMP (C). There was moderate evidence for a shorter AP half-width (D) and increased maximum spike frequency (E) in layer 5 pyramidal neurons relative to layer 2/3.

**Table 8.** Analysis of Effects from Bayesian ANOVA on evoked spike frequency. NOTE: Only effects with BF_incl_>3 have been included in the table.

| Effects | P(incl) | P(excl) | P(incl\|data) | P(excl\|data) | $BF_{incl}$ |
| --- | --- | --- | --- | --- | --- |
| Group | 0.978 | 0.022 | 1.000 | 0.000 | ∞ |
| Sex | 0.978 | 0.022 | 1.000 | $3.414 \times 10^{-7}$ | 65986.434 |
| Hemisphere | 0.978 | 0.022 | 1.000 | $2.324 \times 10^{-8}$ | 969433.124 |
| Layer | 0.978 | 0.022 | 1.000 | $1.204 \times 10^{-9}$ | $1.872 \times 10^{+7}$ |
| Current step (pA) | 0.978 | 0.022 | 1.000 | 0.000 | ∞ |
| Group * Sex | 0.735 | 0.265 | 1.000 | $3.332 \times 10^{-6}$ | 108094.084 |
| Group * Hemisphere | 0.735 | 0.265 | 0.905 | 0.095 | 3.418 |
| Group * Layer | 0.735 | 0.265 | 0.997 | 0.003 | 138.943 |
| Sex * Layer | 0.735 | 0.265 | 0.995 | 0.005 | 66.491 |
| Group * Sex * Hemisphere | 0.265 | 0.735 | 0.856 | 0.144 | 16.496 |
| Group * Sex * Layer | 0.265 | 0.735 | 0.994 | 0.006 | 463.564 |

**Table 9.**
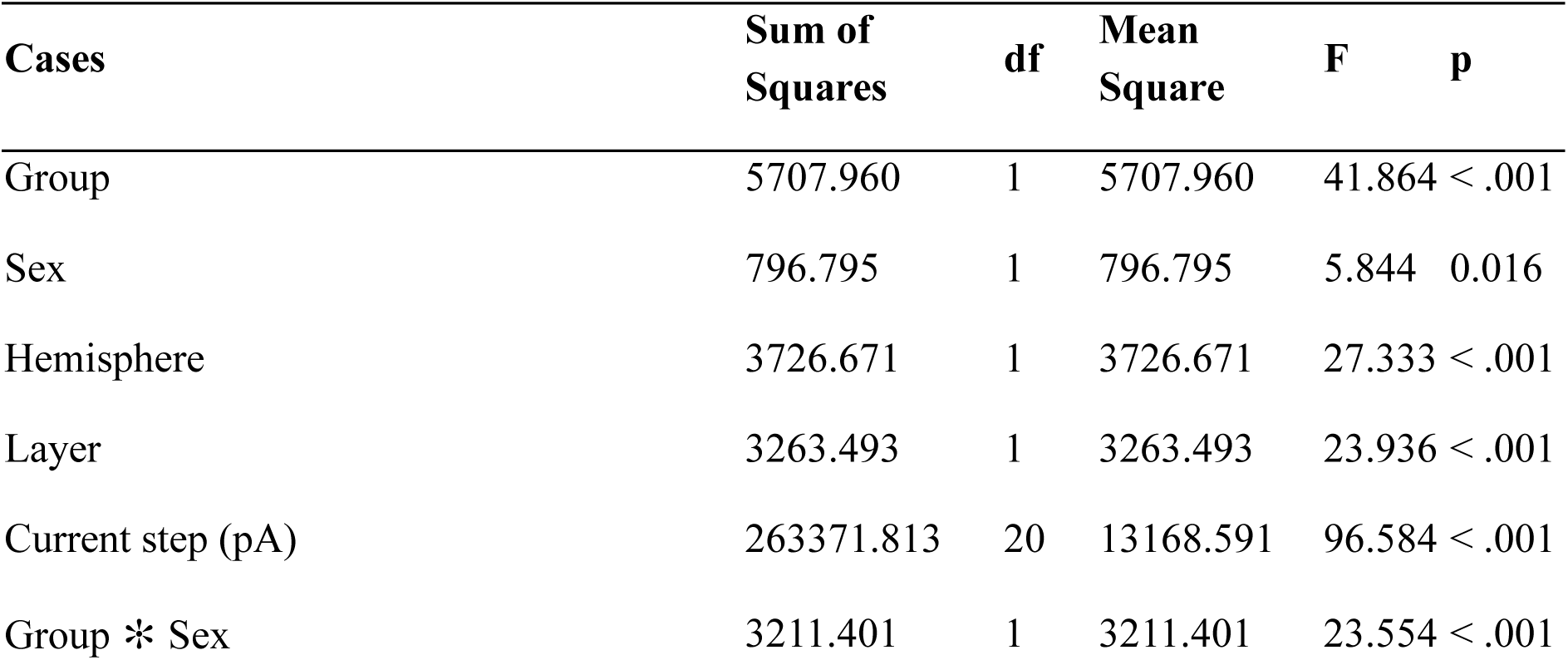
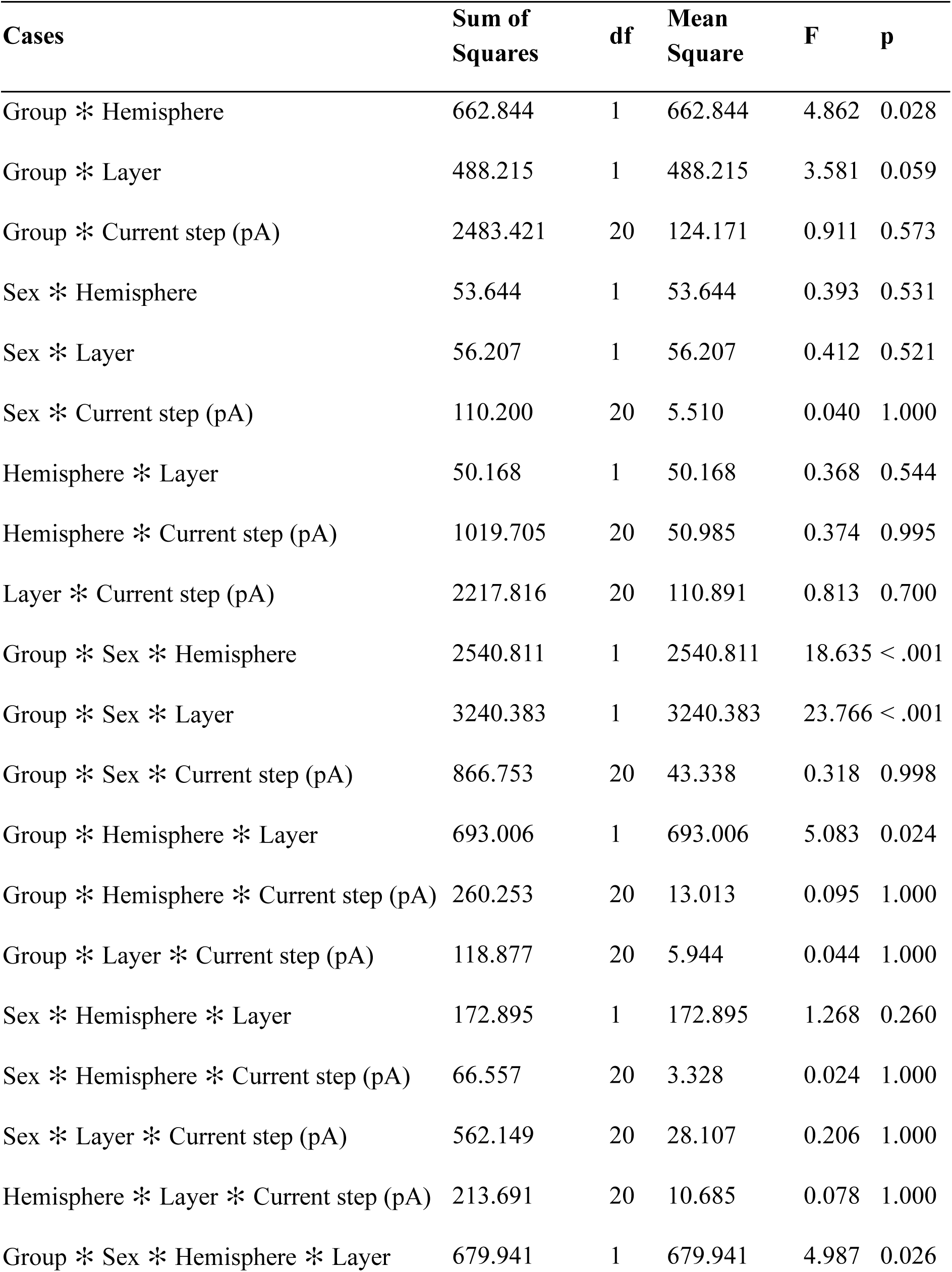
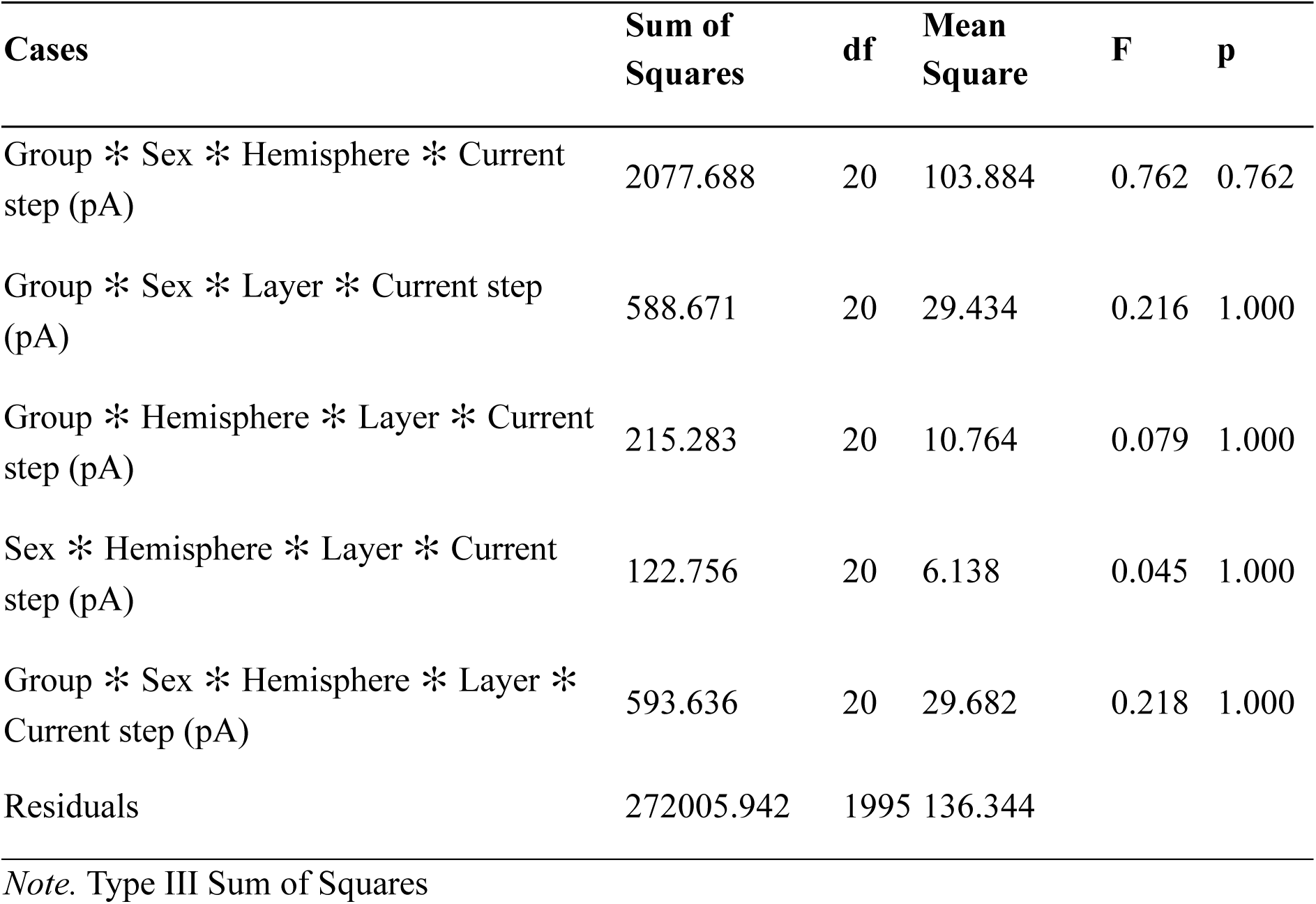
Summary of frequentist ANOVA on evoked spike frequency.

| Cases | Sum of Squares | df | Mean Square | F | p |
| --- | --- | --- | --- | --- | --- |
| Group | 5707.960 | 1 | 5707.960 | 41.864 | < .001 |
| Sex | 796.795 | 1 | 796.795 | 5.844 | 0.016 |
| Hemisphere | 3726.671 | 1 | 3726.671 | 27.333 | < .001 |
| Layer | 3263.493 | 1 | 3263.493 | 23.936 | < .001 |
| Current step (pA) | 263371.813 | 20 | 13168.591 | 96.584 | < .001 |
| Group * Sex | 3211.401 | 1 | 3211.401 | 23.554 | < .001 |

**Table 9. Summary of frequentist ANOVA on evoked spike frequency.**
| **Cases** | **Sum of Squares** | **df** | **Mean Square** | **F** | **p** |
| --- | --- | --- | --- | --- | --- |
| Group * Hemisphere | 662.844 | 1 | 662.844 | 4.862 | 0.028 |
| Group * Layer | 488.215 | 1 | 488.215 | 3.581 | 0.059 |
| Group * Current step (pA) | 2483.421 | 20 | 124.171 | 0.911 | 0.573 |
| Sex * Hemisphere | 53.644 | 1 | 53.644 | 0.393 | 0.531 |
| Sex * Layer | 56.207 | 1 | 56.207 | 0.412 | 0.521 |
| Sex * Current step (pA) | 110.200 | 20 | 5.510 | 0.040 | 1.000 |
| Hemisphere * Layer | 50.168 | 1 | 50.168 | 0.368 | 0.544 |
| Hemisphere * Current step (pA) | 1019.705 | 20 | 50.985 | 0.374 | 0.995 |
| Layer * Current step (pA) | 2217.816 | 20 | 110.891 | 0.813 | 0.700 |
| Group * Sex * Hemisphere | 2540.811 | 1 | 2540.811 | 18.635 | < .001 |
| Group * Sex * Layer | 3240.383 | 1 | 3240.383 | 23.766 | < .001 |
| Group * Sex * Current step (pA) | 866.753 | 20 | 43.338 | 0.318 | 0.998 |
| Group * Hemisphere * Layer | 693.006 | 1 | 693.006 | 5.083 | 0.024 |
| Group * Hemisphere * Current step (pA) | 260.253 | 20 | 13.013 | 0.095 | 1.000 |
| Group * Layer * Current step (pA) | 118.877 | 20 | 5.944 | 0.044 | 1.000 |
| Sex * Hemisphere * Layer | 172.895 | 1 | 172.895 | 1.268 | 0.260 |
| Sex * Hemisphere * Current step (pA) | 66.557 | 20 | 3.328 | 0.024 | 1.000 |
| Sex * Layer * Current step (pA) | 562.149 | 20 | 28.107 | 0.206 | 1.000 |
| Hemisphere * Layer * Current step (pA) | 213.691 | 20 | 10.685 | 0.078 | 1.000 |
| Group * Sex * Hemisphere * Layer | 679.941 | 1 | 679.941 | 4.987 | 0.026 |

**Table 9. Summary of frequentist ANOVA on evoked spike frequency.**
| **Cases** | **Sum of Squares** | **df** | **Mean Square** | **F** | **p** |
| --- | --- | --- | --- | --- | --- |
| Group * Sex * Hemisphere * Current step (pA) | 2077.688 | 20 | 103.884 | 0.762 | 0.762 |
| Group * Sex * Layer * Current step (pA) | 588.671 | 20 | 29.434 | 0.216 | 1.000 |
| Group * Hemisphere * Layer * Current step (pA) | 215.283 | 20 | 10.764 | 0.079 | 1.000 |
| Sex * Hemisphere * Layer * Current step (pA) | 122.756 | 20 | 6.138 | 0.045 | 1.000 |
| Group * Sex * Hemisphere * Layer * Current step (pA) | 593.636 | 20 | 29.682 | 0.218 | 1.000 |
| Residuals | 272005.942 | 1995 | 136.344 |  |  |

*Note.* Type III Sum of Squares

## References

1. Feigin VL, Brainin M, Norrving B, Martins S, Sacco RL, Hacke W, Fisher M, Pandian J, Lindsay P. World Stroke Organization (WSO): Global Stroke Fact Sheet 2022. International Journal of Stroke. 2022;17:18–29.

2. Young BM, Holman EA, Cramer SC, Shah S, Griessenauer CJ, Patel N, Lin DJ, Gee J, Moon J, Schwertfeger J, et al. Rehabilitation Therapy Doses Are Low After Stroke and Predicted by Clinical Factors. Stroke. 2023;54:831–839.

3. Inouye M, Kishi K, Ikeda Y, Takada M, Katoh J, Iwahashi M, Hayakawa M, Ishihara K, Sawamura S, Kazumi T. Prediction of functional outcome after stroke rehabilitation. American Journal of Physical Medicine & Rehabilitation. 2000;79:513–518.

4. Alkayed NJ, Harukuni I, Kimes AS, London ED, Traystman RJ, Hurn PD. Gender-Linked Brain Injury in Experimental Stroke. Stroke. 1998;29:159–166.

5. Reeves MJ, Bushnell CD, Howard G, Gargano JW, Duncan PW, Lynch G, Khatiwoda A, Lisabeth L. Sex differences in stroke: epidemiology, clinical presentation, medical care, and outcomes. The Lancet Neurology. 2008;7:915–926.

6. Thomas Q, Crespy V, Duloquin G, Ndiaye M, Sauvant M, Béjot Y, Giroud M. Stroke in women: When gender matters. Revue Neurologique. 2021;177:881–889.

7. Phan HT, Blizzard CL, Reeves MJ, Thrift AG, Cadilhac DA, Sturm J, Heeley E, Otahal P, Vemmos K, Anderson C. Factors contributing to sex differences in functional outcomes and participation after stroke. Neurology. 2018;90:e1945–e1953.

8. Joy MT, Carmichael ST. Encouraging an excitable brain state: mechanisms of brain repair in stroke. Nature Reviews Neuroscience. 2021;22:38–53.

9. Bütefisch CM, Netz J, Wessling M, Seitz RJ, Hömberg V. Remote changes in cortical excitability after stroke. Brain. 2003;126:470–481.

10. Cicinelli P, Pasqualetti P, Zaccagnini M, Traversa R, Oliveri M, Rossini PM. Interhemispheric asymmetries of motor cortex excitability in the postacute stroke stage: a paired-pulse transcranial magnetic stimulation study. Stroke. 2003;34:2653–2658.

11. Clarkson AN, Huang BS, MacIsaac SE, Mody I, Carmichael ST. Reducing excessive GABA-mediated tonic inhibition promotes functional recovery after stroke. Nature. 2010;468:305–309.

12. Mohajerani MH, Aminoltejari K, Murphy TH. Targeted mini-strokes produce changes in interhemispheric sensory signal processing that are indicative of disinhibition within minutes. Proceedings of the National Academy of Sciences. 2011;108:E183–E191.

13. Boddington LJ, Reynolds JNJ. Targeting interhemispheric inhibition with neuromodulation to enhance stroke rehabilitation. Brain Stimulation. 2017;10:214–222.

14. Imbrosci B, Neitz A, Mittmann T. Focal cortical lesions induce bidirectional changes in the excitability of fast spiking and non fast spiking cortical interneurons. PLoS One. 2014;9:e111105.

15. Paz JT, Christian CA, Parada I, Prince DA, Huguenard JR. Focal cortical infarcts alter intrinsic excitability and synaptic excitation in the reticular thalamic nucleus. Journal of Neuroscience. 2010;30:5465–5479.

16. Grubb MS, Burrone J. Activity-dependent relocation of the axon initial segment fine-tunes neuronal excitability. Nature. 2010;465:1070–1074. doi: 10.1038/nature09160

17. Kuba H, Oichi Y, Ohmori H. Presynaptic activity regulates Na(+) channel distribution at the axon initial segment. Nature. 2010;465:1075–1078. doi: 10.1038/nature09087

18. Schafer DP, Jha S, Liu F, Akella T, McCullough LD, Rasband MN. Disruption of the Axon Initial Segment Cytoskeleton Is a New Mechanism for Neuronal Injury. The Journal of Neuroscience. 2009;29:13242–13254.

19. Hinman JD, Rasband MN, Carmichael ST. Remodeling of the axon initial segment after focal cortical and white matter stroke. Stroke. 2013;44:182–189.

20. Knezic A, Broughton BRS, Widdop RE, McCarthy CA. Optimising the photothrombotic model of stroke in the C57BI/6 and FVB/N strains of mouse. Scientific Reports. 2022;12:7598.

21. Ho J, Tumkaya T, Aryal S, Choi H, Claridge-Chang A. Moving beyond P values: data analysis with estimation graphics. Nature Methods. 2019;16:565–566.

22. Keysers C, Gazzola V, Wagenmakers E-J. Using Bayes factor hypothesis testing in neuroscience to establish evidence of absence. Nature Neuroscience. 2020;23:788–799.

23. Carmichael ST, Archibeque I, Luke L, Nolan T, Momiy J, Li S. Growth-associated gene expression after stroke: evidence for a growth-promoting region in peri-infarct cortex. Experimental Neurology. 2005;193:291–311.

24. Jamann N, Dannehl D, Lehmann N, Wagener R, Thielemann C, Schultz C, Staiger J, Kole MHP, Engelhardt M. Sensory input drives rapid homeostatic scaling of the axon initial segment in mouse barrel cortex. Nature Communications. 2021;12:23.

25. Tang AD, Hong I, Boddington LJ, Garrett AR, Etherington S, Reynolds JN, Rodger J. Low-intensity repetitive magnetic stimulation lowers action potential threshold and increases spike firing in layer 5 pyramidal neurons in vitro. Neuroscience. 2016;335:64–71.

26. King ES, Tang AD. Intrinsic plasticity mechanisms of repetitive transcranial magnetic stimulation. The Neuroscientist. 2024;30:260–274.

27. Beros J, King E, Clarke D, Jaeschke-Angi L, Rodger J, Tang A. Static magnetic stimulation induces structural plasticity at the axon initial segment of inhibitory cortical neurons. Scientific Reports. 2024;14:1479.

28. Neumann-Haefelin T, Witte OW. Periinfarct and remote excitability changes after transient middle cerebral artery occlusion. Journal of Cerebral Blood Flow & Metabolism. 2000;20:45–52.

29. Brown CE, Aminoltejari K, Erb H, Winship IR, Murphy TH. In vivo voltage-sensitive dye imaging in adult mice reveals that somatosensory maps lost to stroke are replaced over weeks by new structural and functional circuits with prolonged modes of activation within both the peri-infarct zone and distant sites. Journal of Neuroscience. 2009;29:1719–1734.

30. Winship IR, Murphy TH. In vivo calcium imaging reveals functional rewiring of single somatosensory neurons after stroke. Journal of Neuroscience. 2008;28:6592–6606.

31. Lim DH, LeDue JM, Mohajerani MH, Murphy TH. Optogenetic mapping after stroke reveals network-wide scaling of functional connections and heterogeneous recovery of the peri-infarct. Journal of Neuroscience. 2014;34:16455–16466.

32. Manwani B, Liu F, Scranton V, Hammond MD, Sansing LH, McCullough LD. Differential effects of aging and sex on stroke induced inflammation across the lifespan. Experimental Neurology. 2013;249:120–131.

33. Hu X, Li P, Guo Y, Wang H, Leak RK, Chen S, Gao Y, Chen J. Microglia/Macrophage Polarization Dynamics Reveal Novel Mechanism of Injury Expansion After Focal Cerebral Ischemia. Stroke. 2012;43:3063–3070.

34. Hurn PD, Macrae IM. Estrogen as a neuroprotectant in stroke. Journal of Cerebral Blood Flow & Metabolism. 2000;20:631–652.

35. Behl C. Oestrogen as a neuroprotective hormone. Nature Reviews Neuroscience. 2002;3:433–442.

36. Choi SG, Shin J, Lee KY, Park H, Kim SI, Yi YY, Kim DW, Song HJ, Shin HJ. PINK1 siRNA-loaded poly (lactic-co-glycolic acid) nanoparticles provide neuroprotection in a mouse model of photothrombosis-induced ischemic stroke. Glia. 2023;71:1294–1310.

37. Keaveney MK, Rahsepar B, Tseng H-a, Fernandez FR, Mount RA, Ta T, White JA, Berg J, Han X. CaMKIIα-positive interneurons identified via a microRNA-based viral gene targeting strategy. Journal of Neuroscience. 2020;40:9576–9588.

38. Veres JM, Andrasi T, Nagy-Pal P, Hajos N. CaMKIIα promoter-controlled circuit manipulations target both pyramidal cells and inhibitory interneurons in cortical networks. Eneuro. 2023;10.

39. Ong RC, Beros JL, Fuller K, Wood FM, Melton PE, Rodger J, Fear MW, Barrett L, Stevenson AW, Tang AD. Non-severe thermal burn injuries induce long-lasting downregulation of gene expression in cortical excitatory neurons and microglia. Frontiers in Molecular Neuroscience. 2024;17:1368905.

